# HMGXB4 Targets *Sleeping Beauty* Transposition to Vertebrate Germinal Stem Cells

**DOI:** 10.1101/2020.06.15.145656

**Authors:** Anantharam Devaraj, Manvendra Singh, Suneel Narayanavari, Guo Yong, Jiaxuan Wang, Jichang Wang, Mareike Becker, Oliver Walisko, Andrea Schorn, Zoltán Cseresznyés, Dawid Grzela, Tamás Raskó, Matthias Selbach, Zoltán Ivics, Zsuzsanna Izsvák

**Affiliations:** Max-Delbrück-Center for Molecular Medicine in the Helmholtz Society (MDC), Robert-Rössle-Strasse 10, 13125 Berlin, Germany; Paul-Ehrlich-Institute, Division of Medical Biotechnology, Paul-Ehrlich-Strasse 51-59, 63225 Langen, Germany

**Author notes:** equal contributions (Anantharam Devaraj, Manvendra Singh). Corresponding authors, Zsuzsanna Izsvák (Z.I), Max-Delbrück-Center for Molecular Medicine (MDC), Robert-Rössle-Str. 10, 13125 Berlin, Germany, Zoltán Ivics.

**Keywords:** *Sleeping Beauty*, transposon, HMGXB4, transcriptional activator, H3K4me3, NuRF complex, chromatin remodelling, nucleolus, SETD1A, SUMOylation, germline, germinal stem cell, Wnt signalling, chromatin domain boundary

## Abstract

Transposons are parasitic genetic elements that frequently hijack key cellular processes of the host. HMGXB4 is a Wnt signalling-associated HMG-box protein, previously identified as a transcriptional regulating host factor of *Sleeping Beauty* (SB) transposition. Here, we establish that HMGXB4 is highly expressed from the zygote stage, and declines after transcriptional genome activation. Nevertheless, HMGXB4 is activated by its own promoter at 4-cell stage, responding to the parental-to-zygotic transition, marks stemness, and maintains its expression during germ cell specification. The HMGXB4 promoter is located at an active chromatin domain boundary. As a vertebrate-specific modulator of SETD1A and NuRF complexes, HMGXB4 links histone H3K4 methyltransferase- and ATP-dependent nucleosome remodelling activities. The expression of HMGXB4 is regulated by the KRAB-ZNF/TRIM28 epigenetic repression machinery. A post-transcriptional modification by SUMOylation diminishes its transcriptional activator function and regulates its nucleolar trafficking. Collectively, HMGXB4 positions SB transposition into an elaborate stem cell-specific transcriptional regulatory mechanism that is active during early embryogenesis and germline development, thereby potentiating heritable transposon insertions in the germline.

## Introduction

HMGXB4 (previously known as HMG2L1) was shown to inhibit Wnt signalling^1^ and smooth muscle differentiation^2^. Nevertheless, HMGXB4 is not commonly recognized as relevant for development.

HMGXB4 was also detected as the most abundant protein in the interactome of the ATP-dependent nucleosome remodelling NuRF (nucleosome remodelling factor) complex^3^, still its role in chromatin remodelling is not characterized. The multi-subunit NuRF complex relaxes condensed chromatin to promote DNA accessibility and transcriptional activation of targeted genes^4–6^. NuRF is a phylogenetically conserved chromatin remodelling complex, originally identified in *Drosophila^7^*. The human core complex of NuRF has similar properties to its *Drosophila* counterpart, and shares the orthologs of three of four components, BPTF (Bromodomain PHD finger transcription factor), SNF2L/SMARCA1 (SWI/SNF Related, Matrix Associated, Actin Dependent Member 1) and the WD repeat containing protein RBAP46/48. The core BPTF contains a PHD finger and a bromodomain that bind to trimethylated histone H3 lysine 4 (H3K4me3) and acetylated histones, respectively^8^.

H3K4me3 is highly enriched at transcription start sites (TSSs) of active genes and controls gene transcription^9,10^. In mammals, SETD1A histone methyltransferase complexes specifically methylate H3K4^11^. SETD1A and NuRF complexes can functionally collaborate to regulate promoter chromatin dynamics (e.g. during erythroid lineage differentiation^12^). Promoters located at the border at Topological Associated chromatin Domains (TADs) are at key genomic positions, offering multiple looping possibilities with neighbouring transcriptional units.

HMGXB4 is a transcriptional activator of the *Sleeping Beauty* transposase^13^. Transposons or transposable elements (TEs) are discrete segments of DNA that have the distinctive ability to move and replicate within genomes across the tree of life. TEs are capable of invading naïve genomes by horizontal transfer (e.g.^14^). The invasion could be successful if the host-encoded factors, required for transposition are phylogenetically conserved, and are readily available in the naïve organism. The general assumption is that TEs (and viruses) piggyback essential host encoded factors to assist their life cycle. HMGXB4 is such a candidate.

*Sleeping Beauty* (SB) was resurrected from inactive transposon copies from various fish genomes^15^. SB transposes via a DNA-based “cut and paste” mechanism, and utilizes several conserved host-encoded factors^13,16–20^. These host-encoded factors regulate transposition throughout the transposition reaction^20^. The SB transposon consists of a single gene encoding the transposase, flanked by two terminal inverted repeats (IRs), which carry recognition motifs for the transposase. The 5’-UTR region of SB can function as a promoter of the transposase^13^, and HMGXB4 enhances transposase expression by interacting with sequences located in the 5’-UTR region of the transposon^13^. SB, by contrast to the Drosophila *P element*, that is controlled by a germline specific splicing process^21^, is not restricted to the germline, and is able to transpose in a wide variety of cells, including both somatic and germinal origin^22–24^.

Nevertheless, since somatic transposition is not heritable, transposons must be targeted to the germ cells to transmit their genome to the next generation. To achieve heritable mobilization, certain TEs transpose in undifferentiated germ cells (primordial germ cells) during embryonic and larval stages and germline stem cells in later developmental stages. Although the host encoded factor targeting factor is unknown, this strategy is used by the *P element* in *Drosophila^25^*. In contrast, retrotransposons barely mobilize directly in germline stem cells^26^. How SB is targeted to the germline is currently unknown.

Following the premise that HMGXB4 is involved in key biological processes, attractive to be captured by a transposon, we used SB transposition as a model to characterize these functions in a developmental context. Our study revealed that HMGXB4 is a vertebrate-specific member of the NuRF complex, and is indeed an essential, but rather overlooked developmental factor, connecting somatic and germinal stemness in early embryogenesis. HMGXB4 targets SB expression to germinal stem cells, where in conjunction with the NuRF complex and SETD1A remodels the chromatin, enhances the expression of the transposase and processes the *de novo* transcripts.

## Results

### HMGXB4 is among the earliest genes expressed in development

As Wnt signalling, which is associated with HMGXB4^1,2^, is one of the earliest cellular processes activated in the developing embryo, we determined the expression profile of HMGXB4 during early development. Our analysis of single cell (sc) transcriptome datasets (scRNA-seq) of mouse and human pre-implantation embryos^27,28^ revealed that HMGXB4 is among the ~300-400 genes detected at significant level (Log2 FPKM > 2) in every cell (Fig. 1a, Supplementary Fig. 1a-b). Furthermore, HMGXB4 is highly expressed prior to embryonic gene activation (EGA), followed by a reduced (but still significant) expression in the preimplantation embryo (Fig. 1a).

**Fig. 1:**
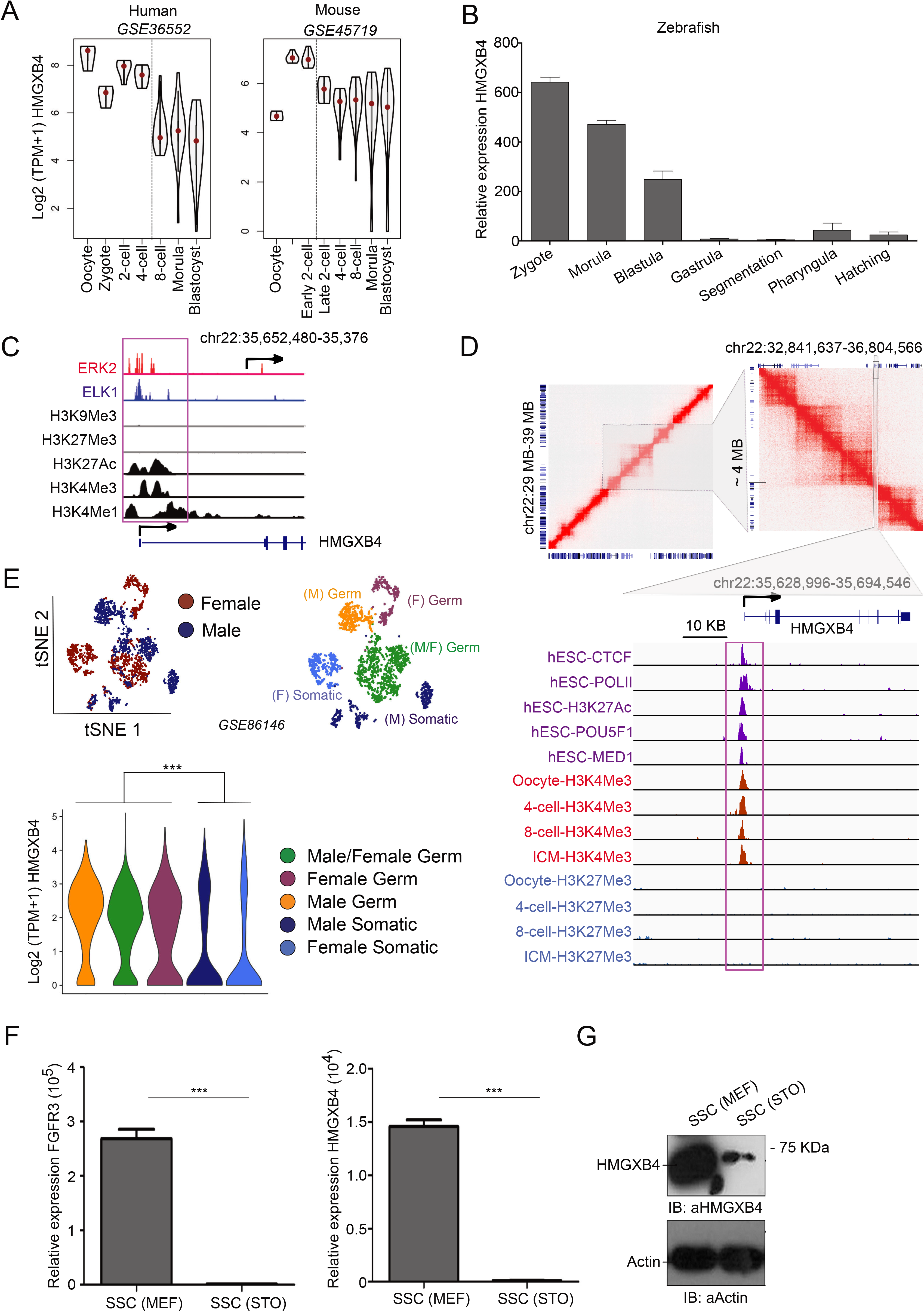
HMGXB4 Connects Pluripotency and Germline. **a** HMGXB4 is expressed before embryonic genome activation (EGA) at the highest level in both human and mouse. Violin plots display the Log2 normalized expression (TPM; Transcript per million) of HMGXB4 in the pre-implantation embryos of human (left) and mouse (right) from the analysis of single cell scRNA-seq datasets. **b** Transcription of HMGXB4 in developing zebrafish embryos (~100) collected at various stages of development, monitored by qRT-PCR (n=2, normalized to GAPDH). **c** HMGXB4 is controlled by a dual occupancy of ERK2/ELK1 transcription factors in pluripotent stem cells. Integrative Genomic Visualization (IGV) of various ChIP-seq raw signals over the HMGXB4 locus in H1-ESCs. Histone modification datasets are from the ENCODE project whereas, ERK2 and ELK1 ChiP-seq datasets are from^31^. **d** Upper panels: Pairwise contact matrices inferred from Hi-C data generated from human embryonic stem cells^29^(GSE116862) show a region between 29MB-39MB on chromosome 22 at 5 kb resolution (left). The intensity of each pixel represents the normalized intensity of observed contacts between a pair of loci. Intensity of red colour is proportional to the intensity of contact between two loci plotted on X and Y axes. Gene models corresponding to these loci are placed on bottom and left side of map. Neighbour loops encompassing ~4MB DNA sequence are further zoomed (right, grey shaded region). HMGXB4 gene is situated at the boundary of the interacting loop (boxed) which is further zoomed (lower panel, grey shaded region): Integrative genome visualization of the active loop boundary around the HMGXB4 locus. Binding profiles of ChIP-seq peaks (boxed) for CTCF, POLII, H3K27Ac, POU5F1 and MED1 (purple peaks) (GSE69646)^63^ over the TSS of HMGXB4 (arrow). CUT&RUN profiling for H3K4me3 (dark red peaks) and H3K27me3 (blue peaks) in human germinal vesicle (GV) oocytes, 4-cell, 8-cell and Inner cell mass (ICM) (GSE124718)^30^ at the TSS of HMGXB4. Note that no significant H3K27me3 peaks were detected on the shown locus. **e** TSNE plot illustrates the clusters of male and female human fetal germ single cells (left panel). The right panel is showing the clusters defined by the top most variable gene expression. The clusters were annotated using published transcriptional markers as male and/or female germ/somatic cells (GSE86146). Violin plots (lower panel) display the Log2 normalized expression of HMGXB4 in the various clusters (colour code is the same as on the middle panel). Every dot represents a single cell. **f** HMGXB4 transcription in spermatogonial stem cells (SSCs) declines upon differentiation (qRT-PCR). Relative expression FGFR3 (left panel) and HMGXB4 (right panel) in rat SSCs cultured on either MEF or STO feeder cells. P value ≤ 0.01. **g** HMGXB4 protein level is reduced in rat SSCs, when cultured on STO feeder cells. Whole-cell lysates of SSC cultured with MEF and STO cells were subjected to immunoblotting using anti-HMGXB4 antibody. (Normalized to GAPDH, P value ≤ 0.01).

In addition to mammalian embryos, we monitored the expression of zHMGXB4 during zebrafish development from the zygote stage to hatching. In conjunction with the mammalian data, our qPCR data shows that zHMGXB4 is highly expressed already in the zebrafish zygote, and its expression level drops from the maternal stage to zygotic transition (Fig. 1b). Following its sharp decline after the blastula stage, zHMGXB4 expression is detectable again in pharyngulas (Fig. 1b). Thus, besides the first steps of embryogenesis, in agreement with its proposed association with Wnt signalling^1,2^, HMGXB4 is expected to act throughout embryonic development.

### HMGXB4 is part of a regulatory network of stemness

To gain insight into the transcriptional regulation of HMGXB4, we performed an integrative analysis of RNA-seq (N ~300), Hi-C, ChiP-seq/ChIP-exo/CUT&RUN of transcription factors (TFs) and histone modification data over the HMGXB4 locus in HeLa, embryonic stem cells H1_ESCs and human early embryogenesis^28–31^. As KRAB-ZNF transcriptional regulators, known to recruit the TRIM28/KAP1-mediated transcriptional repression machinery to specific gene targets in early vertebrate development^32^, we mapped ChIP-exo seq peaks of 230 KRAB-ZNF proteins^33^ around the genomic locus of HMGXB4. We also determined the expression dynamics of the potential regulators during early embryogenesis. Our approach uncovered that HMGXB4 expression is activated by both MAPK1 (alias ERK2) and ELK1 transcription factors (Fig. 1c), and identified repressive KRAB-ZNF proteins (e.g. ZNF468, ZNF763 and ZNF846) harbouring significant peaks (adjusted p-value < 1e-7) at the transcription start site (TSS) of HMGXB4 (Supplementary Fig. 1d-e). Notably, while, the expression of HMGXB4 matches the dynamic of MAPK1/ERK2 throughout the human preimplantation embryogenesis, it is antagonistic to the ZNF468 repressor (Supplementary Fig. 1c). Collectively, our analyses suggest that HMGXB4 is part of a regulatory network of stemness, implicated in coordinating pluripotency and self-renewal pathways^31^, and is expected to be epigenetically controlled by repressive histone marks.

The analysis of 3-Dimensional (3D) conformation of human ESC genomes revealed that the promoter of HMGXB4 is marked by CTCF (Fig. 1d), and co-occupied by ChIP-seq peaks for H3K27ac, MED1 (Mediator 1), POU5F1/OCT4 and POLII over the TSS of HMGXB4, connecting gene expression and chromatin architecture^34^. Adding additional layers of CUT&RUN data analysis for H3K4me3 and H3K27me3 uncovers that the TSS of HMGXB4 has an enrichment for H3K4Me3 but not for H3K27Me3 in human 4-cell, 8-cell and ICM (inner cell mass), indicating that the promoter of HMGXB4 is active in all the stages of pre-implantation development. In addition, the promoter is active in germinal vesicle (GV) stage oocytes, suggesting that the expression of HMGB4 might link somatic and germinal stem cells. The 3D and ChiP analyses suggest that the promoter of HMGXB4 forms a boundary at active compartments (Fig. 1d). This key genomic boundary position might enable multiple interactions between enhancers and promoters in (both somatic and germinal) stem cells.

### HMGXB4 links pluripotent and germinal stem cells

Detecting a high expression signal in oocytes led us to analyse single cell transcriptome datasets of germ cells at several developmental time points (GSE86146) as well as datasets of sex-specific germ cells (GSE63818)^35,36^. We readily observed elevated HMGXB4 expression in both female and male germ cells, compared with somatic cells in the same niche (Fig. 1e and Supplementary Fig. 1f-g). In addition, we analysed scRNA-seq data of germ cells upon differentiation from pluripotent stem cells *in vitro* (GSE102943)^37^. This analysis revealed that HMGXB4 was expressed at comparable levels in both pluripotent and germinal stem cells (CD38^+^), and that the expression of HMGXB4 was maintained during the pluripotent to germ cell transition (Supplementary Fig. 1f). Its expression, by contrast, declined in differentiated germ cells (CD38^−^) (Supplementary Fig. 1f), suggesting that HMGXB4 expression is specific to stem cells. This pattern of HMGXB4 expression is supported by the analysis of a large cohort of scRNA-seq datasets of gonad development (~ 3000 single cells, GSE86146)^35^ (Fig. 1e), identifying HMGXB4 as a novel factor specific for pluripotent and germinal stem cells.

To substantiate the differential expression of HMGXB4 between stem *versus* differentiated germ cells, we used a mammalian (rat) spermatogonial stem cell (SSC) differentiation model. These SSCs maintain their stemness on mouse embryonic fibroblast (MEF) feeders, and differentiate when MEFs are replaced by STO (SNL 76/7) cells^38^. In conjunction with the single cell transcriptome analyses, this approach supported the specific expression of HMGXB4 in spermatogonial stem cells, whereas its expression levels (both transcript and protein) sharply dropped upon differentiation (Fig. 1f-g), indicating that the expression of HMGXB4 is tightly regulated between self-renewing and differentiated states.

### HMGXB4 activates *Sleeping Beauty* transposition in the germline

HMGXB4 has been identified as a host-encoded of factor of SB transposition, serving as a transcriptional activator of transposase expression^13^. Specific expression of HMGXB4 in the germline tempted us to ask whether HMGXB4 is a host-factor that potentiates SB transposition in the germline. To answer, we established a quantitative SB transposon excision assay in SSCs, cultured on MEF or on STO cells (Fig. 2a-b). In the assay, SB transposase expression is driven by the transposon’s 5’UTR containing the sequences at which HMGXB4 transactivates SB transcription. Our assay revealed that the frequency of SB excision was high in SSCs kept on MEFs, while sharply declined upon culturing on STO cells, which triggers differentiation (Fig. 2b). Thus, the rate of transposon excision matches the expression level of HMGXB4, suggesting that HMGXB4 is likely a host-encoded factor associated with activating SB transposition in germinal stem cells. Notably, SB excision, at a decreased level, still occurs in differentiated cells (Fig. 2b), agreeing with the assumption that the requirement of HMGXB4 for transposition is not absolute^13^.

**Fig. 2:**
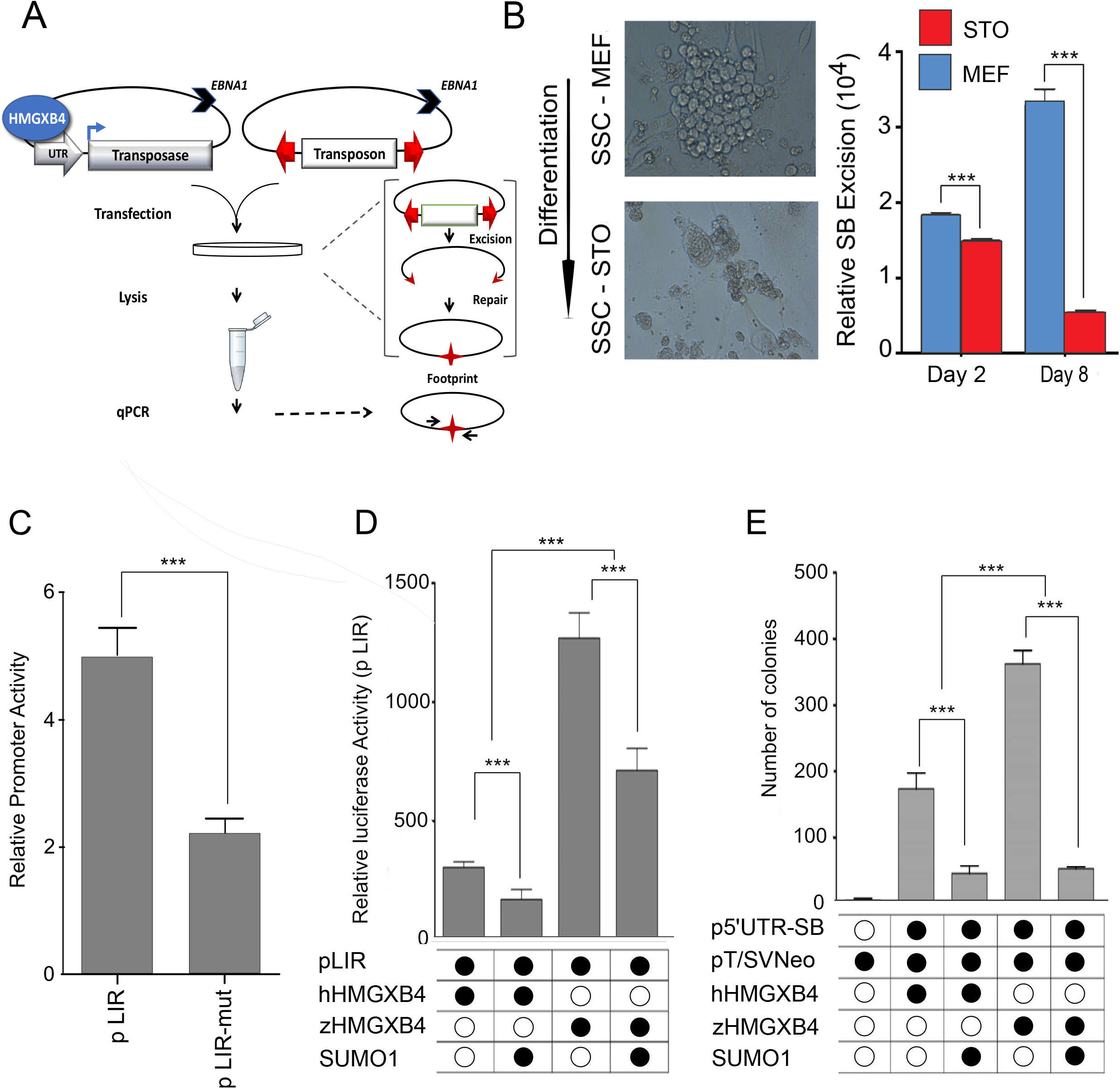
HMGXB4 Targets *Sleeping Beauty* Transposition to the Germline. **a** Schematic of the quantitative transposon excision assay. Both of the plasmid-based transposon and the transposase constructs have an EBNA1 gene providing replication in eukaryotic cells. Transposase expression is driven by the 5’UTR promoter of the SB transposon. HMGXB4 binds the 5’ regulatory region of SB, resulting in an enhanced transposase expression^13^. The transposon is flanked by terminal IRs (red arrows), carrying recognition sequences for the transposase. Following transfection into SSCs, the transposase excises the transposon, leaving a footprint (red star) behind. The footprint is quantifiable using qPCR. **b** (Left panel) Microscopic capture of undifferentiated and differentiated spermatogonial stem cells (SSCs) from the testes of rats. SSC grown on mouse embryonic feeders (MEF) remains undifferentiated, whereas it differentiates by replacing MEF to STO^38^ (8 days). (Scale bar, 40 μm). (Right panel) Excision of the SB transposon declines upon SSC differentiation. The quantitative transposon excision assay (A) detects transposon excision from transiently transfected cells at day 2 and day 8. Continued culturing on MEF (blue) replacing MEF to STO (red). P value ≤ 0.01. **c** The activity of the 5’UTR regulatory region of the SB transposon with/without (wo) the HMGXB4 responding region in zebrafish embryos. One-cell stage zebrafish embryos (~100) were microinjected with a luciferase reporter construct driven by the 5’UTR promoter of SB (located in the left IR, pLIR). A deleted version of the inverted repeat (pΔLIR), not responding to HMGXB4^13^, was used as a control. 36 hours after microinjection, a Dual-Luciferase reporter assay was performed. P value ≤ 0.01. **d** Comparison of the transcription modulating effect of HMGXB4 of either fish or human origin. HeLa cells were transiently co-transfected with a luciferase reporter construct under the control of the SB 5’UTR promoter (located in the left IR, pLIR) and expression plasmids for zHMGXB4 or hHMGXB4 and SUMO1. 48h post-transfection cells were harvested and analysed for luciferase activity. P value ≤ 0.01. **e** Comparison of the effect of HMGXB4 expression of either fish or human origin on SB transposition. HeLa cells were co-transfected with a neo reporter construct, pT/SVNeo, a SB transposase expression construct under the control of SB 5’UTR promoter (located in the left IR, pLIR) and zHMGXB4 or hHMGXB4 in the presence and absence of SUMO1 into HeLa cells, and were subjected to a colony formation (transposition) assay^15^. P value ≤ 0.01

### The transcriptional activation function of HMGXB4 is conserved in vertebrates

A returning question of transposon-host interaction studies concerns their cross-species conservation. While the HMGXB4 gene exists in all vertebrate species, the coding sequence of the fish version is significantly divergent (35%) from its human counterpart (Supplementary Fig. 2a-b). As SB is originated from fish genomes^15^, we asked if the transcriptional enhancer effect^13^ of the human (h)HMGXB4 on SB transposition was reproducible in (zebra)fish embryos. In a reporter assay, luciferase expression was controlled by the 5’-UTR region of the transposon or by a mutated version, where the HMGXB4-responding region was deleted (pLIRΔHRR)^13^. Luciferase activity driven by the 5’-UTR was detectable in zebrafish (*Danio rerio*) embryo extract and depended on the presence of the HMGXB4 responding region (Fig. 2c). In addition, we transiently overexpressed zebrafish (z)HMGXB4 or (h)HMGXB4 by co-injecting the corresponding expression constructs with the reporter into zebrafish embryos. The presence of zHMGXB4 elevated the transcription of the 5’UTR-luciferase reporter three-fold above the level obtained using (h)HMGXB4 in a similar assay (Fig. 2d). The activator effect of zHMGXB4 was even higher compared to the human ortholog when tested in a colony forming transposition assay performed in HeLa cells (Fig. 2e). Collectively, while SB transposition responded more robustly to zHMGXB4 compared to hHMGXB4, the effect/pattern was similar, confirming our hypothesis that HMGXB4 is a conserved host factor of SB transposition in vertebrates^20^. Thus, the transcriptional activation function of HMGXB4 can be modelled by SB transposition from fish to human cells.

### HMGXB4 is post-transcriptionally regulated by SUMOylation

Despite its potential central role in vertebrate embryonic development, the molecular function of HMGXB4 is mostly uncharacterized. Thus, we thought to determine the interacting partners of the HMGXB4 protein. First, we performed a yeast two-hybrid assay (Y2H), using a human HeLa cDNA library (Supplementary Material). The screen identified SUMO1 and PIAS1 (protein inhibitor of activated STAT), suggesting that HMGXB4 is likely modulated post-translationally by the components of the SUMOylation machinery. Co-IP analyses confirmed protein-protein interaction between HMGXB4 and either SUMO1 or PIAS1 in HeLa cells (not shown and Supplementary Fig. 3a).

To find out if HMGXB4 is covalently modified by SUMO1^39^, a tagged HMGXB4-HA (either human or zebrafish origin) was co-expressed with SUMO1, and the protein extracts were analysed by Western blotting (Fig. 3a and Supplementary Fig. 3b). This approach detected a slower migrating band in the presence of SUMO1. Re-probing validated the shifted band to represent SUMOylated protein product (Fig. 3a), suggesting that SUMO1 specifically modifies HMGXB4 via a covalent bond formation to diglycine. To find out if only SUMO1 or other members of the SUMO family, such as SUMO2 and SUMO3 might also modify HMGXB4, expression constructs of the SUMO1,2,3 were co-transfected into HeLa cells, and protein extracts were analysed by Western blotting (Supplementary Fig. 3c). This approach detected slower migrating bands in the presence of all of the tested versions of SUMO, though the most intensive signal appeared (as two shifted bands) in the presence of SUMO1 (Supplementary Fig. 3c). Thus, while all the three SUMO versions could modify HMGXB4, SUMO1 is the most potent modifier.

**Fig. 3:**
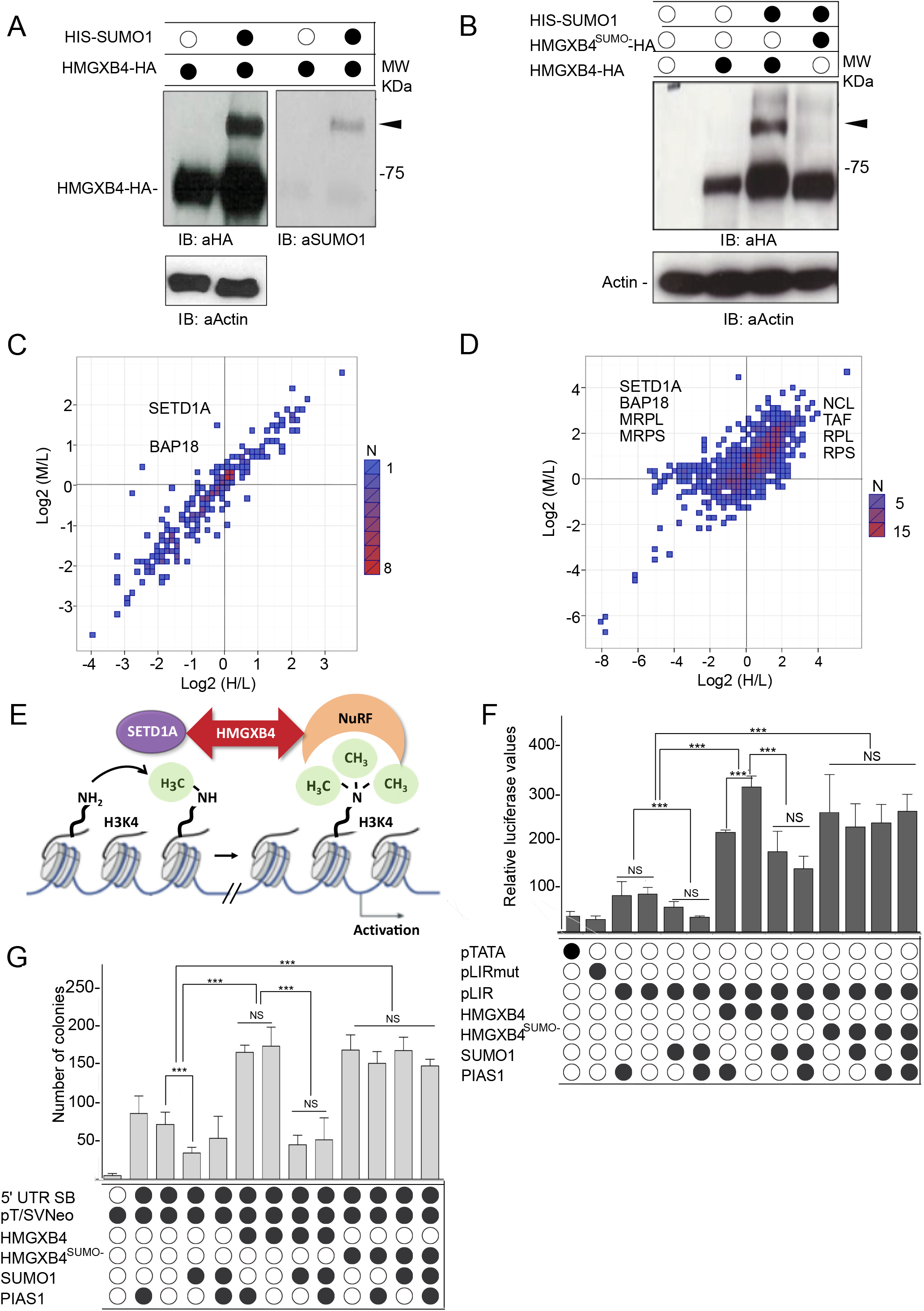
SUMOylation Interferes with the Transcriptional Activator Function of HMGXB4. **a** (Left panel) HMGXB4 gets SUMOylated in the presence of SUMO1 (immunoblots). HeLa cells were co-transfected with expression constructs of the tagged versions of the candidate proteins, HMGXB4-HA and HIS-SUMO1. Whole-cell lysates were immunoblotted. A slower migrating band, potentially corresponding to the SUMOylated version of HMGXB4 is marked by a black triangle (Right panel). The membrane is striped and re-hybridized using an antibody against SUMO1. **b** K317R/K320R dual mutations abolish the post-translational modification, SUMOylation of HMGXB4. HeLa cells were co-transfected with expression plasmids encoding HMGXB4-HA, HMGXB4^K317R/K320R^-HA (referred as HMGXB4^SUMO-^ in the following) and His-SUMO1. The wholecell HeLa lysates were immunoblotted with a HA-specific antibody. The SUMOylated HMGXB4 appears as an additional and slower migrating band (marked by black triangle). **c** SILAC on HMGXB4 pull-down shows the interactome of SUMOylated and non-SUMOylated versions of HMGXB4 from HEK293 cells. Scatter diagram showing the comparison between the two conditions. X-axis represents the Log2-fold change of SUMOylated HMGXB4 vs controls (M/L), whereas Y-axis represents the Log2-fold change of non-SUMOylated HMGXB4 vs controls (H/L). See Fig. S4C for experimental design and the description of Heavy, Medium and Light conditions. Note that SETD1A and BAP18/C17orf49 are detected in the interactome of the non-SUMOylated HMGXB4. **d** The SILAC interactome reveals that in the presence of the SB transposase the interactions are more intense. Scatter diagram showing the comparison between experimental conditions used in SILAC. X-axis represents the Log2-fold change of SUMOylated HMGXB4^WT^ vs controls (M/L), whereas Y-axis represents the Log2-fold change of non-SUMOylated HMGXB4^SUMO-^ vs controls in the presence of the SB transposase (H/L). See Fig. S4C for experimental design and for the description of Heavy (H), Medium (M) and Light (L) conditions. The most significant differentially enriched proteins in the interactomes of SUMOylated HMGXB4^WT^ and of non-SUMOylated HMGXB4^SUMO-^ in the presence of the SB transposase are shown. **e** HMGXB4 provides a link between SETD1A (H3K4Me3 mark deposition) and NuRF (H3K4Me3 reader) complexes to activate the transcription of a set of target genes (schematic model). H3K4Me3 (histone modification for promoters), SETD1A (methyl transferase on H3K4). Note that BPTF (chromatin remodeller, NuRF) and BAP18/C17orf49 (chromatin remodeller around promoters) were identified as interacting partners of HMGXB4^SUMO-^ (see Fig. 3c). **f** SUMOylation interferes with the transcriptional activator function of HMGXB4. The effect of SUMO1 and PIAS1 on SB transposase expression is shown using a Dual Luciferase Reporter assay. Luciferase reporter assays were performed using HeLa cell lysates, where reporter constructs were transiently co-transfected with wild type or mutant HMGXB4 and PIAS1 in various combinations. The luciferase reporter plasmids were either under the control of TATA-box (pTATA) or SB 5’UTR promoter (pLIR LUC). pΔLIR LUC is lacking the HMGXB4 response motif and a minimal promoter (TATA-box) were used as a control. Expression constructs for PIAS1, SUMO1, wild-type HMGXB4 and SUMOylation mutant of HMGXB4^SUMO-^ were used. Standard errors of the mean are from three independent transfections. P value ≤ 0.01. **g** SUMOylation mitigates the transposition activator function of HMGXB4. Effect of SUMO1 and PIAS1 expression on SB transposition. The transposase (driven by its own promoter, 5’UTR SB) and the marker construct pT2/SVNeo are co-transfected with expression constructs of SUMO1 and PIAS1, wild-type HMGXB4 and SUMOylation mutant of HMGXB4^SUMO1-^ in various combinations into HeLa cells and subjected to a colony forming transposition assay. As an additional control, we used a CMV-driven transposase expression construct, not regulated by HMGXB4 (not shown). Standard errors of the mean are from three independent transfections. P value ≤ 0.01.

To map the SUMOylated lysine (K) residues of HMGXB4, we selected those that were phylogenetically conserved among vertebrate orthologs of HMGXB4 (Supplementary Fig. 3d), and converted them to arginine (R) by site-specific mutagenesis (Supplementary Fig. 3e). While most of the K to R substitution mutants only partially affected SUMOylation, the combination of K317R and K320R mutations abolished the two SUMOylated bands (Fig. 3b and Supplementary 3e). We used this version, called as HMGXB4^SUMO-^, for further experiments.

### SUMOylation of HMGXB4 is reversible via SENP-mediated deconjugation, is stress sensitive and does not depend on PIAS1

SUMOylation is a highly dynamic reversible process enabling transient responses to be elicited, which is controlled by conjugating and de-conjugating enzymes. The SUMO moiety can be removed by the SENP [SUMO1/sentrin/SMT3)-specific peptidase] family of SUMO-specific proteases, SENP(1-3) and recycled in a new SUMOylation cycle (reviewed in^40^). To decipher which SENPs de-conjugate SUMO from HMGXB4, we tested SENP1, SENP2 and SENP3 with SUMO1, SUMO2 and HMGXB4-HA in *in vitro* SUMOylation assays. As expected^41,42^, both SENP1 and SENP2 reduced SUMO1 conjugation, whereas SENP3 deconjugated SUMO2 (Fig. 4a-b). Notably, SUMO1 modification of HMGXB4 is sensitive to the presence of the chemical stress factors, ethanol and H_2_O_2_, suggesting a potential additional layer of regulation by stress (Fig. 4c). In these conditions, SB transposition has a slight (~120 %), but reproducible elevation (not shown).

**Fig. 4:**
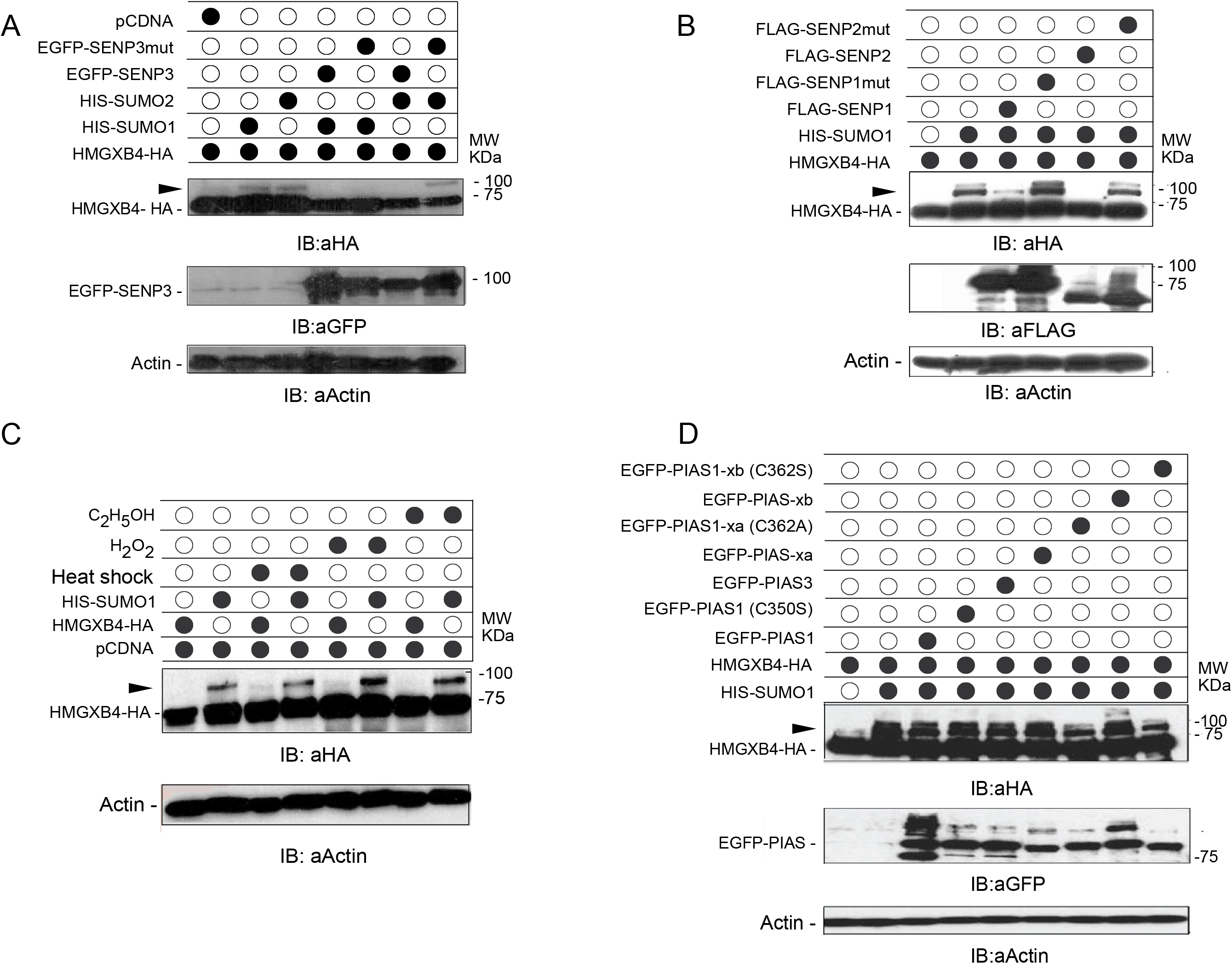
SUMO-specific Proteases (SENPs) and Cellular Stress Affect the SUMOylation of HMGXB4. **a-b** SUMOylation of HMGXB4 is reversible by SUMO-specific proteases (SENPs). HeLa cells were co-transfected with expression constructs of HMGXB4-HA, HIS-SUMO1 and SENP1, SENP2, SENP3 or their SUMO deconjugation defective mutant versions (mut), SENP1^602S^, SENP2^C547S^ and SENP3^C532S42,64^, respectively. Whole-cell lysates were subjected to immunoblotting with anti-HA to monitor HMGXB4, as well as FLAG/GFP–specific antibodies to detect the SENP proteins. The SUMOylated HMGXB4 appears as an additional and slower migrating band (marked by black triangle). **c** SUMOylation of HMGXB4 is stress sensitive. The *in vitro* SUMOylation assay was performed under various cellular stress conditions. Note that ethanol and H_2_O_2_ treatments enhanced the abundance of the SUMOylated bands compared to the untreated condition. The SUMOylated HMGXB4 appears as an additional and slower migrating band (marked by black triangle). **d** PIAS does not affect the SUMOylation of HMGXB4. HeLa cells were co-transfected with expression constructs of HMGXB4-HA, HIS-SUMO1, EGFP-PIAS1, PIAS-xa, PIAS-xa and their respective catalytic mutants in various combinations. Whole-cell lysates were subjected to immunoblotting with anti-HA antibody to detect SUMO modifications, while GFP–specific antibody to detect the PIAS proteins. The SUMOylated HMGXB4 appears as an additional and slower migrating band (marked by black triangle).

In principle, SUMOylation can be also facilitated by E3 ligases, such as PIAS proteins^43^. As PIAS1 was identified as an interactor partner of HMGXB4 in our Y2H assay (confirmed also by co-IP (Supplementary Fig. 3a), we tested various members of the PIAS family, as well as their mutant versions, incapable of SUMO E3 ligase activity (e.g. PIAS1(C350A), PIASxα (C362A) and PIASxβ (C362S)^44^ in a SUMOylation assay (Fig. 4D). Our results argue against a role of PIAS1 as an E3 ligase for HMGXB4. E3 SUMO ligase-independent activities of the HMGXB4-recruited PIAS1 might involve transcriptional coregulation^45^ (not followed up in the current study).

### Non-SUMOylated HMGXB4 links histone H3K4 Methyltransferase- and ATP-dependent nucleosome remodelling activities

SUMOylation might affect several aspects of the target protein, including structure, interaction partners, cellular localization, enzymatic activity or stability (reviewed in^40^). HMGXB4 has an estimated half-life of ~30 hours (Protparam/Expasy). Notably, both HMGXB4^wt^ and HMGXB4^SUMO-^ were detectable at similar levels at different timepoints following a cyclohexamide treatment (Supplementary Fig. 4a-b), indicating that SUMOylation had no effect on the stability of HMGXB4 protein. Alternative to a hypothesis-driven strategy, we have performed an unbiased high throughput protein interactome analysis to decipher the effect of SUMOylation on HBGXB4 function. We used a triple SILAC pull-down approach, suitable for relative quantification of proteins by mass spectrometry^46^. We also included the SB transposase in the assay in order to find out which functions of HMGXB4 are affected by the presence of the transposase. Thus, in the experimental setup, we transfected HEK-293T cells with HA-tagged HMGXB4^wt^ and HMGXB4^SUMO-^ in the presence/absence of SB transposase and/or SUMO1 (Supplementary Fig. 4c).

According to gene ontology (GO) analysis, the interactome of HMGXB4 could be characterized by the top terms of *Wnt signalling, Ribonucleoprotein complex, Structural and cytoskeletal, Host response to viruses, Transcriptional regulation, Translation initiation, Nonsense-mediated decay (NMD*) and *Regulation of metabolism* (Fig.s S5A-B). In the presence of the SB transposase, the top GO categories were not changed (Supplementary Fig. 5c), but the affinity of the interactions became stronger and/or the number of interacting partners became higher in most of the shared GO categories (Fig. 3c-d and Supplementary Fig. 5d), suggesting that the transposase modulates these functions of HMGXB4 and intensifies interactions to its partners.

While the interactomes of HMGXB4^wt^ and HMGXB4^SUMO-^ were highly related, SUMOylation specifically affected the affinity of HMGXB4 to its interaction partners in two groups of proteins. Firstly, in the HMGXB4^SUMO-^ interactome, we detected the BAP18 (BPTF associated protein of 18 kDa, alias (*C17orf49*) (Fig. 3c and Supplementary Fig. 5e). BAP18, in association with the ATP-dependent NuRF active (H3K4me3) chromatin reader complex^3^, has been previously reported in androgen receptor induced transactivation^47^. In addition, among the most differentially recruited proteins of HMGXB4^SUMO-^, we also identify SETD1A (Fig. 3c-d and Supplementary Fig. 5f), a histone-Lysine N-methyltransferase generating mono-, di- and trimethylation at H3K4 (H3K4me1-3) at transcriptional start sites of target genes^48^, indicating that HMGXB4 links members of a multiprotein complex that participates in both depositing and reading active chromatin marks at H3K4 (Fig. 3e). Notably, in comparison to HMGXB4^SUMO-^, the wildtype HMGXB4 has a lower affinity for both SETD1A and C17orf49/BAP18 (Fig. 3c-d), suggesting that the activation of target gene expression by HMGXB4 is controlled by SUMOylation. Thus, non-SUMOylated HMGXB4 links histone H3K4 methyltransferase- and ATP-dependent nucleosome remodelling activities.

To confirm the effect of SUMOylation on the transcriptional activator function of HMGXB4, using the SB model, we performed both transcription and transposition assays in the presence of either HMGXB4^SUMO-^ or HMGXB4^WT^. In the transient luciferase reporter assay, 5’UTR-luciferase, HMGXB4^wt^, HMGXB4^SUMO-^, SUMO1 and PIAS1 were co-transfected into HeLa cells in various combinations. In agreement with the observation that non-SUMOylated version of HMGXB4^SUMO-^ interacts with H3K4Me3 (via C17orf49/BAP18 and SETD1A), the presence of SUMO1 attenuated the transcriptional activation function of HMGXB4 (Fig. 3f). Accordingly, the HMGXB4-mediated enhancement on transposition^13^ was also diminished in the presence of co-transfected SUMO1 (Fig. 3g). Similar to its human ortholog, SUMO1 modulated the activity of zHMGXB4 in both assays (Fig. 2d-e). Collectively, these data support that SUMOylation diminishes the transcriptional activator function of HMGXB4. In line with the results of the *in vitro* SUMOylation experiments (Fig. 4d), elevated PIAS1 levels had no obvious consequence on SB activity in either of the assays (Fig. 3f-g).

### SUMOylation induces nucleolar compartmentalization of HMGXB4

The second notable group of the differential proteome of HMGXB4^wt^/HMGXB4^SUMO-^ was associated with nucleolar functions (> 25%) (Fig. 5A), suggesting that the SUMOylation might affect nucleolar compartmentalization and activities. The nucleolar interactors had higher affinity to HMGXB4^WT^, and were associated *Translation initiation and elongation, Transcriptional control* (Fig. 5b), *Nonsense-mediated decay, Ribonucleoprotein complex (ribosomal structure*) (Fig.s S5A-B). The presence of the SB transposase intensified the affinity of interaction in all of these GO categories (Fig. 3d and Supplementary Fig. 5d). Thus, via HMGXB4, the transposase might sponge on transcription activation, non-sense-mediated decay, transcript processing and protein translation machineries of the host cell.

**Fig. 5:**
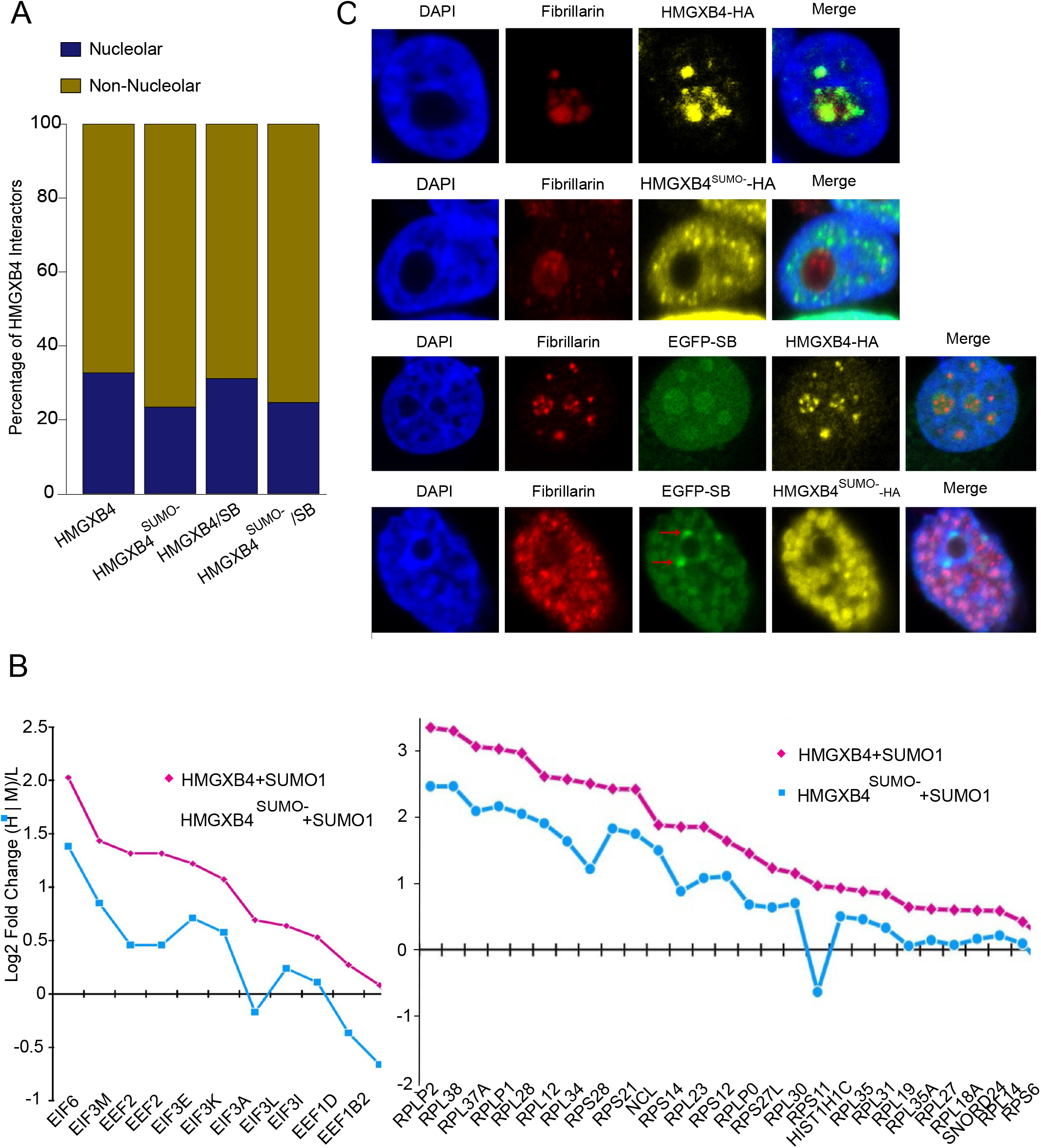
SUMOylated HMGXB4 is Compartmentalised to the Nucleolus. **a** SUMOylation affects the affinity of HMGXB4 to its nucleolar interacting partners. The stacked bar plot visualizes the differential affinity of HMGXB4 and HMGXB4^SUMO-^ with nucleolus associated proteins in the presence and absence of the SB transposase. **b** Line plots display the effect of SUMOylation on HMGXB4 to a selected set of protein interacting partners. Proteins involved in *Translation initiation and elongation* (left panel); in *Transcriptional control* (right panel). Note that the depicted proteins are localized in nucleolus (experimental evidence). **c** Visualization of sub-nuclear localization of transiently expressed HMGXB4 or HMGXB4^SUMO-^, using immuno-fluorescent confocal microscopy in HeLa cells. Upper two rows; HMGXB4-HA (yellow); fibrillarin (red); DAPI (blue); merge; Lower two rows; as on the upper panels, but in the presence of transiently co-expressed SB transposase (EGFP-SB, green). Note that the SB transposase co-localizes with both wildtype HMGXB4 and HMGXB4^SUMO-^, except in the perinuclear nuage (red arrows). Curiously, the coexpression of HMGXB4^SUMO-^ and SB transposase disrupts the integrity of the nucleolus, which mobilizes the fibrillarin marker all over the cytoplasm. (Scale bar, 40 μm).

To validate nucleolar localization and its regulation by SUMOylation, we used confocal microscopy to monitor subcellular trafficking of HMGXB4 upon SUMOylation. We co-transfected expression vectors of HA-tagged HMGXB4, HMGXB4^SUMO-^, EGFP-tagged SUMO1 (EGFP-SUMO1) and SB (EGFP-SB) into HeLa cells in various combinations, and subjected the cells to microscopy. This strategy revealed an antagonistic subcellular localization pattern of HMGXB4 and HMGXB4^SUMO-^ (Fig. 5c and Supplementary Fig. 6), supporting the prediction of our differential interactome data. While HMGXB4^SUMO-^ stayed in the nucleoplasm, HMGXB4 co-localized with the nucleolar marker fibrillarin, confirming that SUMOylation regulates the subnuclear trafficking of HMGXB4 (Fig. 5c and Supplementary Fig. 6), and thus physical sequestration from the NuRF/SETD1A complex. It is notable that while, endogenous SUMO1 level is capable of supporting the nucleolar trafficking of HMGXB4, HMGXB4^SUMO-^, in the presence of SB transposase partially disrupts the integrity of the nucleolus, which mobilizes the fibrillarin marker all over the cytoplasm (Fig. 5c).

SB transposase co-localized with either or with HMGXB4^SUMO-^ in the nucleoplasm or HMGXB4 in the nucleolus (Fig. 4c and Supplementary Fig. 6), suggesting that SB transposition piggybacks both nuclear and nucleolar functions of HMGXB4, involved in transcription initiation and transcript processing, respectively. Curiously, unlike HMGXB4, the SB transposase is enriched in the perinuclear nuage of cells (Fig. 4c), where the machinery of piRNA biogenesis is concentrated^49^.

## Discussion

Viruses and transposons frequently piggyback ‘essential’ cellular mechanism(s) of the host. The role of HMGXB4 in *Sleeping Beauty* (SB) transposition is conserved from fish to human, supporting the assumption that HMGXB4-SB transposon interaction can be generally modelled in vertebrates^20^. Here, we used the HMGXB4-SB host-parasite interaction model to decipher certain cellular function(s) of the transposon-targeted, but otherwise poorly characterized developmental gene, HMGXB4. Our study identifies HMGXB4 as a novel factor linking pluripotency to the germline, and the host encoded factor that shepherds SB transposition to germinal stem cells.

HMGXB4 is among the first expressed genes in the embryo, and in agreement with its regulatory role in Wnt signalling^1,2^, its expression level is dynamically changing throughout embryogenesis. Following maternal expression, HMGXB4 is activated by its own promoter at 4-cell stage, responding to the parental-to-zygotic transition. HMGXB4 marks stemness, and maintains its expression during germ cell specification. The promoter of HMGXB4 is located at an active chromatin domain boundary in stem cells, potentially offering multiple looping possibilities with neighbouring genomic regions. Thus, beside the germline, the recruitment of HMGXB4 supports efficient SB transposition during early embryogenesis in various somatic progenitor cells, suggesting that HMGXB4 is primarily recruited as a spatio-temporal transcriptional activator of the transposase in stem and progenitor cells.

HMGXB4 provides a physical bridge between BAP18 and SETD1A, thereby linking histone H3K4 methyltransferase- and ATP-dependent nucleosome remodelling activities. Notably, HMGXB4 is not conserved outside vertebrates, thus it provides a vertebrate-specific function(s) to the core NuRF complex, first identified in *Drosophila^7^*.

Via HMGXB4, SB piggybacks a multiprotein complex, capable of both depositing and reading active chromatin marks at H3K4. Furthermore, via ERK2/MAPK1-ELK1, MED1, CTCF and POU5F1, HMGXB4 is part of the transcription regulatory network, implicated in pluripotency and self-renewal^31^.

HBGXB4 is regulated by a reversible post-translational modification, SUMOylation. While SUMOylation does not affect the stability of the HMGXB4 protein, it regulates its binding affinity to its protein interacting partners. The non-SUMOylated HMGXB4 recruits the SETD1A/NuRF complex, and acts as a transcriptional activator, whereas SUMOylation serves as a signal for its nucleolar partition. The recruitment of HMGXB4 from the nucleoplasm to the nucleolus provides a flexible regulation of the transcription activating epigenetic machinery by affecting the stoichiometry of the HMGXB4 containing protein complexes.

The SB transposase follows its host factor during its subnuclear trafficking to the nucleolus, thus piggybacks both SUMOylated and non-SUMOylated functions of HMGXB4. In addition to its well-characterized role in ribosome biogenesis^50^, the nucleolus is involved in several other crucial functions, including maturation and assembly of ribonucleoprotein complexes, cell cycle regulation and cellular aging. These nucleolar functions are frequently targeted by several viruses to support their own replication (reviewed in^51^). For example, regulatory viral proteins, such as the accessory protein 3b from the SARS-CoV, affecting cell division and apoptosis, predominantly localizes in the nucleolus^52,53^.

Interestingly, the SB transposase, but not HMGXB4, is enriched in the perinuclear nuage-like structure, associated with piRNAs, known to repress transposable elements via RNAi (reviewed in^49^). Curiously, SB is not endogenous in human, thus there are no SB-specific piRNAs present in human cells, suggesting that SB might be capable of recognizing an evolutionary conserved feature of Piwi-interacting small RNA (piRNA) biogenesis.

The HMGXB4-mediated germline targeting is a likely conserved feature of the *Tc1-like* family of transposons (where SB belongs) in vertebrates. In addition to the *Tc1-like* elements, the *Drosophila P* element utilizes a similar strategy to target germinal stem cells^25^. While it has been shown that the targeting process in *Drosophila* is controlled by the piRNA pathway^25^, the host encoded targeting factor of *P* element transposition is yet to be identified, and could not be identical to the vertebrate specific HMGXB4.

Unlike retrotransposons that rarely mobilize in undifferentiated germinal stem cells^26^, SB directly targets this cell type. Retrotransposons, by contrast, use an indirect approach. In this scenario, certain *Drosophila* retrotransposons were shown to “hijack” the microtubule transporting system to transfer their transcripts from the interconnecting supporting nurse cells to the transcriptionally inactive oocyte^26^. Mammalian retrotransposons likely use a similar scheme^54^. The more aggressive, direct targeting strategy used by DNA transposons (e.g. *Sleeping Beauty, P element*) is expected to generate a higher level of germline toxicity, and might – at least partially – explain the evolutionary success of retrotransposons over DNA transposons in higher vertebrates.

As a third strategy, TEs were suggested to manipulate the blastomere to adopt a germinal, rather than somatic fate. During this process, TE-derived sequences have been incorporated into gene regulatory networks of the pluripotent cells^19,55–58^.

While, HMGXB4 is involved in epigenetic regulation of gene expression itself, it is controlled by the stress-sensitive KRAB-ZNF/TRIM28-mediated epigenetic repression mechanism^59^. In addition, the SUMO-specific conjugation of HMGXB4 is also a stress-inducible, dynamic process, and thus could activate transcription upon environmental changes. The stress-sensitiveness of HMGXB4 would enable SB transposon to sense and react to cellular stress, a known feature of transposable elements^60^. Similar, a stress responsive SUMO-regulated chromatin modification has been also implicated in reactivating integrated viruses in the genome (e.g. heterochromatin histone demethylase, JMJD2A in Kaposi’s sarcoma associated herpes virus (KSVH)^61^.

Importantly, our current work on deciphering a relationship between a host-encoded factor piggybacked by a transposable element also spotlights on so far overlooked aspects of HMGXB4. Besides nucleosome remodelling, HMGXB4 is involved in modulating downstream regulatory processes of target gene activation and production. The activity of HMGXB4 is stem/progenitor cell specific, and the expression level of HMGXB4 drops sharply upon differentiation and stays at an undetectable level in differentiated cells. Nevertheless, HMGXB4 is epigenetically regulated, stress sensitive and when expressed, it can support target gene activation in any cell type. Thus, aberrant activation of HMGXB4 in differentiated cells (e.g. cancer) might result in undesirable gene expression. In this context, it is notable that HMGXB4 has been identified as a target of epithelial splicing regulatory proteins upon epithelial-mesenchymal transition (EMT)^62^, suggesting that HMGXB4 could be an important target in future cancer research.

## Supporting information

HMG Supplemental Table

HMG Supplemental Figure 1

HMG Supplemental Figure 2

HMG Supplemental Figure 3

HMG Supplemental Figure 4

HMG Supplemental Figure 5

HMG Supplemental Figure 6

## Supplementary Information

### Supplementary Figures

**Supplementary Fig. 1. HMGXB4 Connects Pluripotency and Germline.**

**(a)** Boxplots showing the expression distribution of 413 genes that are highly expressed (Log2 FPKM > 2) in every single cell (> 99%) of human preimplantation embryos (Supplementary Table 2). Besides known housekeeping genes (e.g. GAPDH and actin, not shown), HMGXB4 is expressed in every cell. **(b)** Top Gene Ontology (GO) of the 413 ubiquitously expressed genes. GO identification numbers from top to bottom. GO:0008135, GO:0003743, GO: 0022857, GO:0003735, GO:0016887, GO:0000184, GO:0004129, GO:0061024, GO:0050793, GO:0007010, GO:0006754, GO:0006885. **(c)** Similar (e.g. activation) and antagonistic (e.g. repression) expression of ERK2/MAPK1 and ZNF468 with HMGXB4. Violin plots display the Log2 normalized expression of ERK2/MAPK1 and ZNF468 in the preimplantation embryos of human from the analysis of single cell RNA-seq datasets. Note that the expression of ZNF468 showed anti-correlation with HMGXB4 expression. EGA, embryonic gene activation. **(d)** Histogram illustrates the distance of 230 KRAB-ZNF ChIP-exo peaks from the Transcription Start Site (TSS) of HMGXB4. ZNF468, ZNF763 and ZNF846 (red) peaks intersected on HMGXB4 TSS. Note that only the expression of ZNF468 showed a robust anticorrelation with HMGXB4 expression (see on Supplementary Fig. 1c), thus predicts ZNF468 as a repressor of HMGXB4. **(e)** KRAB-ZNF468 binding interferes with the transcription of HMGXB4. Line plot shows the density of KRAB-ZNF468 raw ChIP-exo signal over the TSS of HMGXB4. See also Supplementary Fig. 1c-d. **(f)** HMGXB4 is expressed throughout the differentiation of human induced pluripotent stem cells (iPSCs) to CD38^positive^ primordial germ (hPG)-like cells, marked by CD38 (GSE102943). Barplots showing the transcriptomic changes of HMGXB4 during the differentiation (Diff) process. Note the downregulation of HMGXB4 in CD38^minus^ (somatic) cells. **(g)** TSNE plot illustrates the single cell clusters from the development of male and female human Primordial Germ (hPG)-like cells (GSE63818) on the basis of the expression of Most Variable Genes (MVGs). The single-cell clusters are distinguishable as Male/Female Germ/Somatic cells. (left panel). Violin plots on the right panel display the Log2 normalized expression of HMGXB4 in the clusters using the same colour codes. Every dot represents a single cell. Note the germ stem cell specific expression of HMGXB4.

**Supplementary Fig. 2. Phylogenetic Conservation of HMGXB4 in Vertebrates.**

**(a)** Conversation of the HMGXB4 sequences in vertebrates. UCSC snapshot displaying the Multiz alignment of HMGXB4 UTRs and coding sequences across the 100 vertebrates at single base pair resolution. Black colour denotes the sequence similarity. Gapped sequences are probable deletions when compared to the human version of HMGXB4. **(b)** Phylogenetic analysis of HMGXB4 proteins across different vertebrate species. Sequences of from various vertebrate species were obtained from NCBI. The sequences were aligned with ClustalW and the tree was generated using Maximum Parsimony.

**Supplementary Fig. 3. HMGXB4 is Regulated by Post-translational Modification, SUMOylation (Related to Fig. 3)**

**(a)** Physical interaction between HMGXB4 and PIAS1. Co-immunoprecipitation of PIAS1 and HMGXB4 in HeLa cell lysates (HMGXB4-HA, PIAS1-FLAG). **(b)** HMGXB4 of zebrafish origin (z) gets SUMOylated in the presence of SUMO1 (immunoblots). HeLa cells were co-transfected with expression constructs of the tagged versions of the candidate proteins, zHMGXB4-HA and HIS-SUMO1. Whole-cell lysates were immunoblotted. A slower migrating band, potentially corresponding to the SUMOylated version of HMGXB4 is marked by a black triangle. **(c)** All the three SUMO variants (e.g. 1, 2 and 3) can SUMOylate HMGXB4. HeLa cells were co-transfected with expression plasmids encoding HMGXB4-HA, HIS-SUMO1, 2, 3 or a mutated version of FLAG–SUMO1ΔGG, defective in conjugation. The SUMOylated HMGXB4 versions are expected to appear as slower migrating bands (by a black triangle) on the immunoblot, using HA-specific antibody in whole-cell lysates. Importantly, no shifted bands were detectable when HeLa cells were transfected with the mutated version of SUMO1(mut), lacking the diglycine C-terminal motif required for its conjugation to substrates SUMO1-ΔGG^44^. **(d)** Sequence alignment to predict phylogenetically conserved Lysin (K) residues as potential SUMOylation sites in the HMGXB4. The position of the mutated K residues that abolish SUMOylation in HMGXB4 marked in red. The double mutant K317R/K320R is referred as HMGXB4^SUMO-^ in further studies. **(e)** Functional testing of putative SUMOylation mutants. Various mutant versions of HMGXB4 were co-transfected with HIS-SUMO1 into HeLa cells and subjected to immunoblotting using a HA tag-specific antibody. A slower migrating band, potentially corresponding to the SUMOylated version of HMGXB4 is marked by a black triangle.

**Supplementary Fig. 4. The Effect of SUMOylation on HMGXB4 Function. SUMOylation does not affect the stability of the HMGXB4 protein (Related to Fig. 3)**

**(a-b)** SUMOylation does not affect the stability of the HMGXB4 protein. Comparison of steady-state levels of HMGXB4 and HMGXB4^SUMO-^ in the presence of SUMO1, (A) untreated and (B) under conditions when *de novo* protein synthesis was blocked by cyclohexamide (inhibitor of protein biosynthesis) treatment. HeLa cells were transfected with HA tagged HMGXB4 or SUMOylation mutant HMGXB4^SUMO-^ along with SUMO1 in triplicates. Cells were lysed 12, 36 or 72 hours posttransfection without or following cycloheximide (inhibitor of protein biosynthesis) treatment. 10 μg of protein were immunoblotted and hybridized with anti-HA antibody. A slower migrating band, potentially corresponding to the SUMOylated version of HMGXB4 is marked by black triangles. **(c)** Triple SILAC pull-down experimental design to investigate the effect of SUMOylation on the interaction partners of HMGXB4 in the absence (Exp 1) or the presence (Exp 2) of the non-hyperactive *Sleeping Beauty* transposase (SB10)^15^. Expression constructs were transiently transfected to HEK293T cells. In the SILAC/pull-down experimental approach, stable isotope labelled amino acids (Light (L) or Medium heavy (M)) are added in the form of medium supplement to culture HEK293T cells. Detection of interaction partners is performed by mass spectrometry (MS).

**Supplementary Fig. 5. Characterization of the HMGXB4^WT^ and HMGXB4^SUMO-^ Interactomes (Related to Fig. 3)**

**(a)** Heatmap displaying the similar and distinct patterns of HMGXB4^WT^ and HMGXB4^SUMO-^ interactomes (SILAC-MS). **(b)** Characterization of HMGXB4 interacting protein partners by Gene Ontology (GO) terms. NMD, nonsense mediated decay. Connected to Supplementary Fig. 5a. **(c)** Venn diagram shows the shared and unique Gene Ontology (GO) terms of the HMGXB4 interacting partners in the presence and absence of SB. **(d)** Bar plot showing the shared GO terms (Fig. S5B) HMGXB4 interacting partners in the presence or absence of SB at the level of – Log10 adjusted-p-value (higher the bar, lower is the false discovery rate). Number of the analysed proteins (N) are indicated next to the colour legends. Total proteins detected in SILAC experiments was kept as a background. Note the more intense interactome in the presence of the SB transposase in all GO categories. **(e)** Co-immunoprecipitation to validate endogenous HMGXB4 and BAP18/C17orf49 interaction in HEK293 cells. **(f)** Co-immunoprecipitation to validate endogenous HMGXB4 and SETD1A interaction in HEK293 cells (predicted by SILAC).

**Supplementary Fig. 6. SUMOylated HMGXB4 is Compartmentalised to the Nucleolus.**

(Upper two rows) Visualization of the sub-nuclear localization of transiently co-expressed EGFP-SUMO1, HMGXB4 or HMGXB4^SUMO-^ using immuno-fluorescent confocal microscopy in HeLa cells. EGFP-SUMO1 (green), HMGXB4-HA (HA-yellow); DAPI (blue), fibrillarin (red); merge. Note the inverse localization of HMGXB4 and HMGXB4^SUMO-^. The similar pattern of HMGXB4 or HMGXB4^SUMO-^ in the presence or absence (Fig. 4c) of co-transfected SUMO1 suggest that excess of SUMO1 did not change the trafficking behaviour of either versions of HMGXB4, suggesting that the endogenous SUMO1 is sufficiently marking HMGXB4^WT^ for the nucleolar transit. Curiously, the expression of HMGXB4^SUMO-^ disrupts the integrity of the nucleolus, which mobilizes the fibrillarin marker all over the cytoplasm. (Lower three rows) Cellular localisation of EGFP, EGFP-SB and EGFP-SUMO1. Note the nucleolar accumulation of SB transposase. (Scale bars 40 μm).

**Supplementary Table 1.**
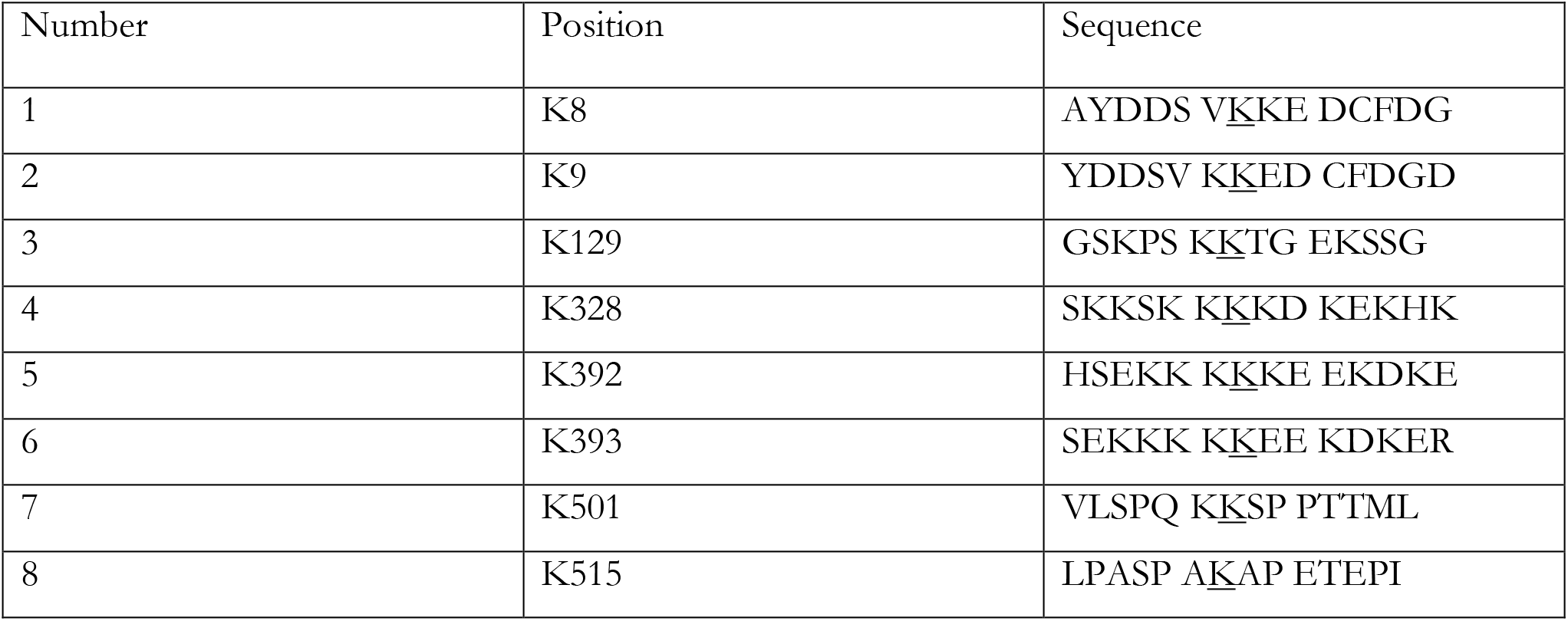
*In silico* predicted SUMOylation sites using (Sumoplot™) www.abgent.com/tools/toSumoplot)^65,66^. The underlined lysine (K) residues were subjected to site directed mutagenesis to arginine (R) and tested in the *in vitro* SUMOylation assay (co-transfecting them with SUMO1 into HeLa cells and subjected to immunoblotting) (Supplementary Fig. 3d).

**Supplementary Table 2**

List of ubiquitously expressed gene in early human development (.xlsx)

## Methods

### Constructs

pCAG-Venus-SB10, The SB10 transposase^15^ gene was cloned into pCAG-Venus, zHMGXB4 coding sequence was PCR amplified from the cDNA of zebrafish embryo with zHMG EcoRI Fwd. and NotI Rev. primers. The PCR product was digested with EcoRI and NotI restriction enzymes and subcloned into the corresponding sites of the pcDNA3.1 vector; HMGXB4-HA, C-terminal HA (hemagglutinin peptide, YPYDVPDYA)-tagged versions of the HMGXB4 were obtained by inserting HA tag downstream of the coding region of the zebrafish/human HMGXB4 protein by PCR, and cloning into the HindIII/XhoI sites of pcDNA3.1.

### Mitotic inactivation of MEFs

Mouse embryonic fibroblasts (MEFs) are often used as feeder cells in embryonic stem cell research. MEFs were isolated from 12.5 to 13.5 post coitum (p.c.) mouse embryos. The embryos were dissociated and then trypsinized to produce single-cell suspensions. After expansion, confluent MEFs cells were treated with 10μg/ml mitomycin-C (Sigma) for 2 hours in DMEM at 37°C. Cells were then washed twice with PBS followed by trypsinization, and counted before dilution and plating.

### Rat spermatogonial stem cells culturing

Rat spermatogonial stem cell lines were cultured on mitomycin-C treated MEFs in spermatogonial culture medium (SG medium) as described^67^. The cells were passaged at 1:3 dilutions onto a fresh monolayer of MEFs every 10–14 days at 3 × 10^4^ cells/cm^2^. For passaging, cultures were first harvested by gently pipetting them free from the MEFs. After harvesting, the clusters of spermatogonia were dissociated by gentle trituration with 20–30 strokes through a p1000 pipette in their SG culture medium. The dissociated cells were pelleted at 200 × g for 4 min, and the number of cells recovered during each passage was determined by counting them on a hemocytometer. Feeder Removal Micro Beads (Miltenyi Biotec) were used for depletion of MEFs while co-culturing with rat spermatogonial stem cells for gene expression and protein studies. The protocol for co-culturing the rat spermatogonial stem cells with STO cells was done as in^38^. The separation of STO cells from rat spermatogonial stem cells was carried using the Magnetic activated cell sorting (MACS) from Miltenyi Biotec.

### Microinjection of zebrafish embryos

In order to measure the promoter activity of the left inverted repeat (LIR) of Sleeping Beauty transposon in zebrafish embryos, a microinjection mix containing 50 ng/μl of firefly luciferase and 0.2 ng/μl Renilla reporter plasmids containing Buffer Tango 10× to a final concentration of 0.5x, and phenol red solution to a final concentration of 0.05% were injected into fertilized eggs of wild type *Danio rerio*. The injected embryos were incubated for 24–48 h at 28.5°C in egg water. For measurement of promoter activity, the egg water is removed and the embryos were washed with PBS followed by lysis with 50 μl of passive lysis buffer (PLB) 1x for 30 min at room temperature, shaking at 150 rpm. Firefly and Renilla luciferase activities were measured according to the manufacturer protocol (Dual-Luciferase Reporter Assay System, Promega).

### Transient expression of proteins in HeLa or HEK293 cells

HeLa or HEK293 EBNA cells were cultured in DMEM medium supplemented with L-glutamine, penicillin/streptomycin and 10% FBS (Invitrogen). Cells were transiently transfected at 50% – 65% confluency with QIAGEN-purified plasmid DNA using JetPEI or FuGENE transfection reagent according to the manufacturer’s instructions. After 48 hours post transfection cells were lysed in RIPA lysis buffer containing 25 mM Tris-HCl pH 7.6, 150mM NaCl, 1% NP-40, 1% sodium deoxycholate, 0.1% SDS supplemented with protease cocktail inhibitors (Roche) and subjected to western blot analysis.

### Total protein quantification

Protein concentrations were measured by using calorimetric technique at a wavelength of 562 nm (OD562) by bicinchoninic acid assay (Pierce™ BCA™ Protein-Assay-Thermo Fisher). Samples containing known concentrations of bovine serum albumin (BSA) were used as a standard.

### SDS-polyacrylamide gel electrophoresis

Proteins were separated by their molecular weight using SDS-polyacrylamide (SDS-PAGE) gels ranging 10-15 %. The total protein (50 μg) lysate (prepared with modified RIPA buffer) was mixed with 1x Laemmli buffer incubated for 5 min at 95°C and resolved at 80 V in SDS-PAGE running buffer (50 mM Tris-HCl, 196 mM glycine, 0.1% SDS, pH 8.4). After electrophoresis the gels were subjected to western blotting.

### Western blotting

Cell or tissue extracts were resolved by 10-15% SDS-polyacrylamide gel electrophoresis (SDS-PAGE) and electro-transferred to polyvinylidene difluoride (PVDF) membranes (Amersham, Inc.). The membranes were incubated with 5% nonfat dry milk at room temperature for 1 hour and probed overnight with specific antibodies at 4°C. The immune complexes were detected by enhanced chemiluminescence (ECL, Pierce) with anti-mouse, anti-goat, anti-rat or anti-rabbit immunoglobulin G (IgG) coupled horseradish peroxidase as the secondary antibody (Pierce).

### Co-immunoprecipitation

HeLa (8.8 x 10^6^) or HEK293 (20 x 10^6^) cells were co-transfected with indicated plasmids. Whole-cell extract was prepared using extraction buffer (Tris-HCl 50 mM at pH 7.4, NaCl 150 mM, EDTA 1 mM, NP-40 1% and Na-deoxycholate 0.25 %) supplemented with protease inhibitor cocktail (Roche, Mannheim, Germany). For immunoprecipitations, equal amounts of lysate (containing 5 mg of total cellular protein from HeLa cells) were pre-cleared with protein G-agarose beads (Sigma, St Louis, MO). Pre-cleared extracts were incubated with 1 μg rat monoclonal anti-HA (Roche,) for 2 h at 4 °C. Precipitates were washed extensively in lysis buffer, bound complexes were eluted with 2x SDS–PAGE sample buffer and resolved by 7.5–10% SDS–PAGE. Immunoblotting was performed according to standard procedures and proteins detected with the indicated antibodies. Antibodies were detected by chemiluminescence using ECL Advance Western Blotting Detection Kit (Amersham Bioscience).

### SUMOylation assay

HeLa cells (8.8×10^6^) were co-transfected with the expression constructs in 10 cm dishes with 2 μg each of His-SUMO1 and HA-tagged wild-type HMGXB4 or mutants. At 48 h post transfection, cells were lysed by radioimmunoprecipitation assay (RIPA) buffer containing 10 mM N-ethylmaleimide (NEM) and protease inhibitors. For immunoprecipitation, equal amounts of lysates (containing 5 mg of total cellular protein from HeLa cells) were pre-cleared with protein G-agarose beads (Sigma, St Louis, MO). Pre-cleared extracts were incubated with 1 μg rat monoclonal anti-HA (Roche, Mannheim, Germany) for 2 h at 4 °C. Precipitates were washed extensively in lysis buffer, bound complexes were eluted with 2x SDS–PAGE sample buffer and resolved by 7.5–10% SDS–PAGE. Immunoblotting was performed according to standard procedures and proteins detected with the anti-HA antibodies. Antibodies were detected by chemiluminescence using ECL Advance Western Blotting Detection Kit (Amersham Bioscience).

### Protein stability assay

HeLa cells (8.8×10^6^) were co-transfected with the expression constructs in 10 cm dishes with 5 μg each of His-SUMO1 and HA-tagged wild-type HMGXB4 or mutants. After 12 h transfection, cells were incubated with 100 μM cyclohexamide (Sigma) for 0 to 72 hour and then harvested with radioimmunoprecipitation assay (RIPA) buffer containing 10 mM N-ethylmaleimide (NEM) and protease inhibitors. Equal amounts of total proteins from each treatment were taken to perform western blot analysis.

### Stable isotope labelling with amino acids in cell culture (SILAC)

This protocol relies on the incorporation of amino acids containing substituted stable isotopic nuclei (e.g. ^12^C, ^13^C and ^13^C/^15^N) into proteins in living cells. The three cell populations are grown in culture media that are identical except that one medium contains a “Light,” and the other two medium a “Medium Heavy” (or Medium) and “Heavy,” form of a particular amino acid (^12^C-Arginine, ^13^C-Arginine and ^15^N-Arginine, respectively). The mass spectra data were analysed using MetaCore from GeneGo Inc (www.genego.com). A fold change cutoff of 0.5 with a p-value < 0.05, was set to identify proteins whose expression was significantly differentially regulated. Enrichment analysis was conducted using GeneGo curated ontologies along with Gene Ontology to provide a quantitative analysis of the most relevant biological functions represented by the data.

### Mass spectrometry

A triple SILAC pull-down experiment using anti-HA resin to investigate interaction partners of HMGXB4 and HMGXB4 defective of SUMOylation in the presence/absence of Sleeping Beauty in transiently transfected HEK293T cells. The cells were cultured in SILAC DMEM, High Glucose (4.5 g/l), w/o L-Arg, L-Lys, L-Gln (PAA), 10% dialyzed fetal bovine serum (PAA),4 mM L-glutamine (PAA),1% penicillin/streptomycin (100 IU/100 μg, Invitrogen). For Light Medium, they were supplemented with 28 mg/l L-arginine ^12^C_6_·^14^N_4_·HCl (Sigma), 48.7 mg/l L-lysine ^12^C_6_·^14^N_2_·HCl (Sigma), Medium Heavy 28 mg/l L-arginine ^13^C_6_·^14^N_4_·HCl (Sigma), 48.7 mg/l L-lysine ^13^C_6_·^14^N_2_·4,4,5,5-D_4_-HCl (Sigma) and Heavy Medium 28 mg/l L-arginine ^13^C_6_·^15^N_4_·HCl (Sigma),48.7 mg/l L-lysine ^13^C_6_·^15^N_2_·HCl (Sigma). The cells were cultured for two doublings and were transiently transfected at 80% confluency with 5μg each of pCMV His-SUMO1, pCMV-SB10 and pIRES HA-tagged wild-type HMGXB4 or mutants using JetPEI transfection reagent according to the manufacturer’s instructions. After 24 h transfection, the cells were lysed in lysis buffer containing 25 mM Tris/HCl, pH 7.4 (Carl Roth),125 mM KCl (Merck),1 mM MgCl2 (Merck),1 mM EGTA/KOH pH 8.0 (Carl Roth),5% glycerol (Merck),1% NP-40 (Nonidet P 40 Substitute, Sigma), 1 mM DTT (Sigma, added freshly) and 1X Protease inhibitor cocktail (Complete, EDTA-free, Roche, added freshly) followed by immunoprecipitation with Anti-HA Agarose beads (Sigma-Aldrich EZview Red Anti-HA Affinity Gel). The precipitated protein complex was detected for interaction partners by mass spectrometry and the results obtained were analyzed by MaxQuant computational platform.

### Quantitative assay to monitor *Sleeping Beauty* transposon excision

Both the transposon and transposase expression constructs were EBNA1-based in order to avoid fast degradation of the plasmids. pEBNA-SB100X and pEBNA-Tneo are based on pCEP4/pEBNA vector (a kind gift from T. Willnow, MDC). For pEBNA–Tneo construct, pEBNA was digested with SnaB1 and Xho1. The Tneo insert was released from the pTNeo construct by Xho1 and Sal1. For pEBNA – SB100X construct, pEBNA was cleaved by BamH1 and Xho1 and ligated to the insert of UTR-SB100X released with BglII and XhoI from pcDNA3.1 UTR SB100X. In pEBNA – SB100X, the transposase is driven by its own promoter located in the left inverted repeat of the transposon (UTR). Following co-transfection of the transposon excision monitoring system (500 ng of each pEBNA-SB100X and pEBNA-Tneo) into SSCs cultured either on MEFs or STO cells, the cells were lysed at the indicated time points. The qPCR detects the precise excision product^16^, generated upon SB excision.

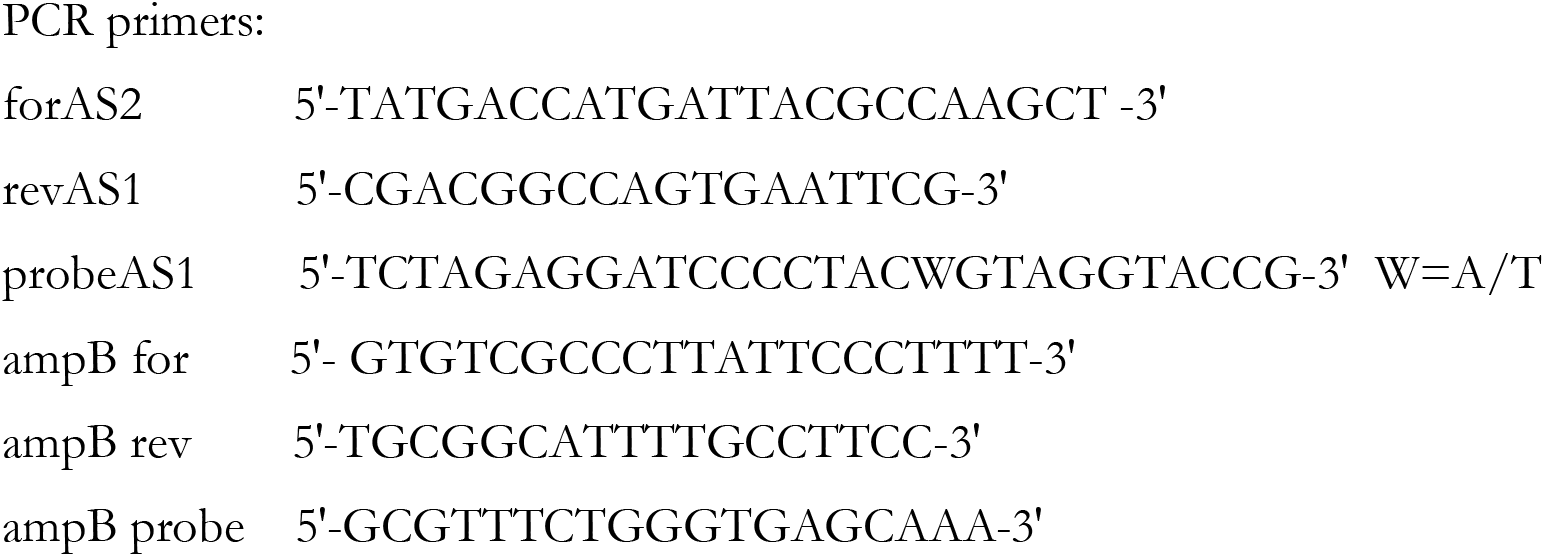

Note: For the ambiguous base in the footprint, a probe was ordered with a degenerate nucleotide in that position to pick up both variations in our PCR.

The Taqman-excision PCR was performed from cell lysates. 3 μl of the lysate was mixed with primers and probes detecting the excision product and a gene on the input plasmid in a total reaction volume of 20 μl using *TaqMan Universal PCR* Master Mix (Applied Biosystems) on the *7900HT Sequence Detection System* (Applied Biosystems). Average Ct values were calculated from quadruplicates of each sample. The amounts of the excision product and input plasmid were calculated using a standard curve for each primer/probe set carried along with each measurement. Standard curves resulted from a dilution row of excision product (pXS plasmid DNA cloned for that purpose) and pEBNA-Tneo input plasmid over a 4-5 logarithmic range. The calculated absolute amounts of excision product were normalized to those of the input plasmid pEBNA-Tneo, and the standard deviation was determined.

Details of an excised pTneo (=pXS) primer/probe binding sites: ACACAGGAAACAGC<u>TATGACCATGATTACGCCAAGCT</u>TGCATGCCTGCAGGTCGAC*TCTAGAGGATCCCCTACWGTAGGTACCG*AGCT<u>CGAATTCACTGGCCGTCG</u>TTTTACAACGTCGTGACTGGGAAAACCCTGGCGTTACCCAACTTAATCGCCTTGC; *underlined: for and rev primer; italic: FAM/TAMRA-probe covering footprint with W = A or T*

### Transposition assay

5 x 10^5^ HeLa cells seeded in 6-well plates were co-transfected with 90 ng of neo^R^ carrying transposon plasmid plus 150 ng of transposase expression plasmid or mock control along with 90 ng each of His-SUMO1, Flag-PIAS1, HA-tagged wild-type HMGXB4 and HA-tagged HMGXB4 defective of SUMOylation. Two days post-transfection, the cells were trypsinized and 1/10 of the cells were seeded on 10 cm-diameter dishes and selection was started by culturing the cells in DMEM supplemented with 600 mg/ml G418 (Biochrom). After 14 days, selection was terminated by washing the cells with phosphate buffered saline (PBS), fixed in 10% v/v formaldehyde and stained with methylene blue in PBS, and counted. The experiments were done at least thrice and results are presented as means plus standard deviations.

### Dual Luciferase reporter assay

The Luciferase reporter assay was carried out to understand the effect of HMGXB4 on 5’UTR promoter activity of SB in the presence and absence of SUMO1 and PIAS1 proteins. 5 x 10^5^ HeLa cells seeded in 6-well plates were co-transfected with 200 ng of luciferase reporter plasmid along with 150 ng each of His-SUMO1, Flag-PIAS1, HA-tagged wild-type HMGXB4 and HA-tagged HMGXB4 defective of SUMOylation. In all samples, 10 ng of the plasmid pRL-TK (Promega) encoding Renilla luciferase were included for normalization of transfection efficiency. After 48 h, cells were lysed and assayed by using the Dual Luciferase kit (Promega). Relative luciferase activity is the ratio of Firefly to Renilla luciferase activity, normalized to the activity of the reporter alone. The experiments were done at least thrice, with at least duplicate samples in each study. Results are presented as means plus standard deviations.

### Immunofluorescence staining

Monolayers of HeLa (0.3 x 10^6^) or cells was grown on coverslips were co-transfected with the expression constructs with 250 ng each of EGFP-SB10, PML-YFP, EGFP-SUMO1, HMGXB4^SUMO-^ -HA and HMGXB4-HA tag using FuGENE transfection reagent according to the manufacturer instructions. Cells were fixed at 36 h post transfection with cold 1% paraformaldehyde in Phosphate-buffered saline (PBS) for 20 min at 4°C and then permeabilized with 1% Tween in PBS and incubated for 2h at room temperature. Cells were then stained for HMGXB4-HA and HMGXB4^SUMO^ -HA using a HA tag antibody (Roche Applied Science) and Fibrillin with Anti-Fibrillin 1 antibody (abcam) for 2 h in a humid chamber at 37°C. Covers slips were then washed 3x with PBS and stained with Anti-Rat IgG conjugated (Novus Biologicals) with Cy7, Anti-Rabbit IgG (abcam) with Alexa Fluor 647 and DAPIfor 1 h at 37°C. The coverslips were removed from the well and rinsed with dH2O to remove excess PBS. Coverslips were placed in Fluorescent Mounting Medium (DAKO Cytomation) on a glass microscope slide and dried overnight. The staining was analysed by confocal microscope (LSM 710 with software ZEN lite 2011; Carl Zeiss, Inc.).

### FACS analysis

Fractions of GFP positive cells were determined by fluorescence-activated cell sorting, using the FACS Calibur (Becton Dickinson). The data was analysed using CELLQuest v. 3.1 (Becton Dickinson). Briefly, cells were trypsinized in 10 cm tissue culture dishes, the reaction was stopped by adding DMEM medium (Gibco) containing 10% FCS. Cells were collected in polystyrene tubes and centrifuged at 2500 rpm, 3 minutes at 4°C and washed with ice cold PBS two times. 50,000 cells were analysed per sample with the same cell flow rates.

### SENP assay

To protect SUMO-conjugated proteins from deSUMOylation, the cell lysates were treated with N-ethylmaleimide (NEM), an inhibitor of SUMO-specific isopeptidases and subjected to Westernblotting.

### SILAC data analysis

We downloaded the list of proteins associated with the nucleolus (supported by experimental evidence) from human proteome atlas (https://www.proteinatlas.org/humanproteome/cell/nucleoli) and intersected with the interactome of HMGXB4 in presence and absence of SB. A Fold change cutoff of 0.5 with a P-value<0.05 was set to identify proteins interacting with HMGXB4. Enrichment analysis was conducted using GeneGo curated ontologies along with Gene Ontology to provide a quantitative analysis of the most relevant biological or molecular functions represented by the data.

### Single cell RNA-seq analysis

Single cell (sc)RNA-seq datasets were downloaded in a raw format from five independent studies: Human embryogenesis (GSE36552), mouse embryogenesis (GSE45719), differentiation of pluripotent stem cells to human primordial germ like cells (hPGC-like) (GSE102943), hPGCs in a given space and time (GSE86146) and development of human germline cells in a gonadal niche (GSE86146). The datasets comprised of ~ 3500 single cellular transcriptomes. The transcription of genes in every cell was calculated at TPM or FPKM expression levels. Samples were included in the analysis only if they had gene expression data of at least 5000 genes with expression levels exceeding the defined threshold (Log2 TPM > 1). We considered only those genes for the analysis that were expressed in at least 1% of the total samples. We used Seurat 1.2.1 from R to normalize the datasets at logarithmic scale using *“scale.factor = 10000”*. After normalization, we calculated scaled expression (z-scores for each gene) for downstream dimension reduction. The cells were separated by subjecting the MVGs ({Log(Variance) & Log2(Average Expression)} > 2) to the dimension reduction methods of principal component analysis (PCA) which were further subjected to TSNE analysis. Consequently, HMGXB4 expression from normalized data in each cluster was calculated and visualized using the same tool.

### Data mining of ChIP-seq datasets

ChIP-seq datasets for histone modifications were obtained from the ENCODE project. ERK2/MAPK1/ELK1 ChIP-seq data were obtained from^31^. H3K27ac, MED1, POU5F1, POLII and CTCF ChIP-seq datasets were obtained from (GSE69646)^63^. Hi-C data was obtained from^29^ (GSE116862) in “.hic” format. These contact matrix files were further normalized and visualized as a heatmap, using “Juicer” tool. ChIP-seq datasets in raw fastq format were aligned against hg19 reference genome by Bowtie (version 2.2.2) under the parameters *“local-sensitive”*. All unmapped reads, non-uniquely mapped reads and PCR duplicates were removed from the analysis. Aligned reads were converted into bedGraph format using genomeCoverageBed from BedTools to visualize utilizing IGV over RefSeq genes (hg19).

### Data mining of ChIP-exo peaks of KRAB-ZNF proteins

ChIP-exo peaks of 230 KRAB-ZNF protein were obtained from GSE78099. Signals from ChIP-exo data were obtained from MACS2 (as in^33^) after running over with parameter *-g hs -q 0.01 -B*. To visualize the overlapping peaks at the HMGXB4 locus, the obtained signals were merged beneath Integrative genome visualizer (IGV) tracks. Density of MACS2 signals were plotted around the TSS of HMGXB4 to show the significant occupancy.

### Statistics

Statistical analyses (Student’s t-test) were performed using the Prism 5 software (GraphPad) for molecular biology experiments. R was used for the statistical analysis of high throughput data. The level of significance was calculated using Wilcoxon-test and corrected by multiple testing.

## Contributions

Planned experiment: Zs.I. and A.D. Performed experiments: A.D. S. N. G.Y. D.G. T. R. J.W. J.W. M.B. Methodology: O.W. A.S. Z.C. M.S. Analysed data: M.S. Writing original draft: Zs.I. Edited final version: Z.I.

