## Supplementary material for "HMGXB4 Targets *Sleeping Beauty* Transposition to Vertebrate Germinal Stem Cells": HMG Supplemental Figure 1

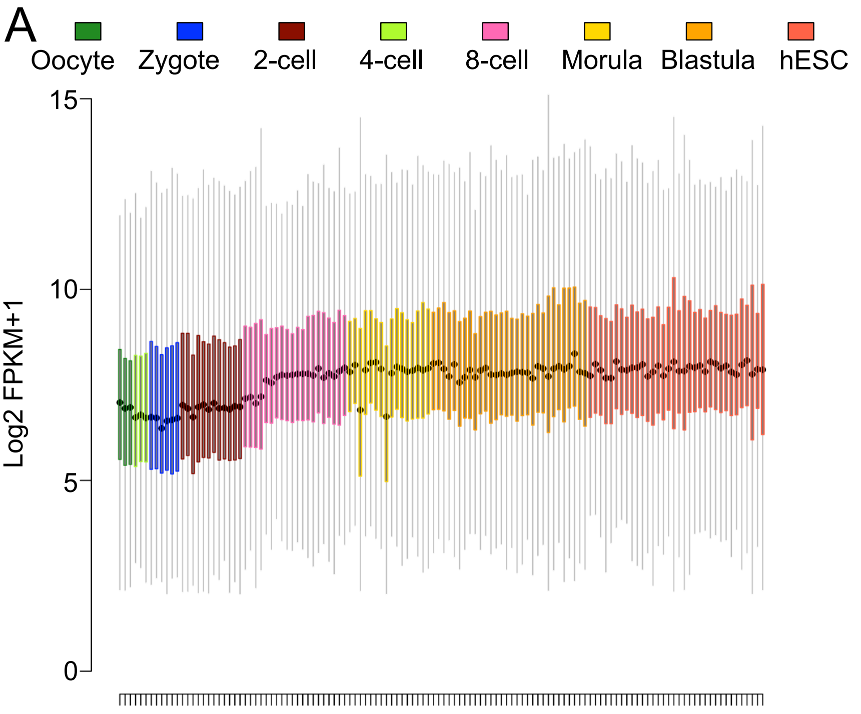


B

| Gene Ontology enrichment | FDR |
|---|---|
| translation factor activity, RNA binding | 4.10E-13 |
| translation initiation factor activity | 7.98E-9 |
| transmembrane transporter activity | 7.70E-8 |
| structural constituent of ribosome | 2.71E-6 |
| ATPase activity | 3.52E-6 |
| mRNA catabolic process | 1.84E-5 |
| cytochrome-c oxidase activity | 3.73E-4 |
| membrane organization | 6.20E-4 |
| regulation of developmental process | 1.92E-3 |
| cytoskeleton organization | 3.33E-3 |
| ATP biosynthetic process | 5.11E-3 |
| regulation of pH | 5.13E-3 |

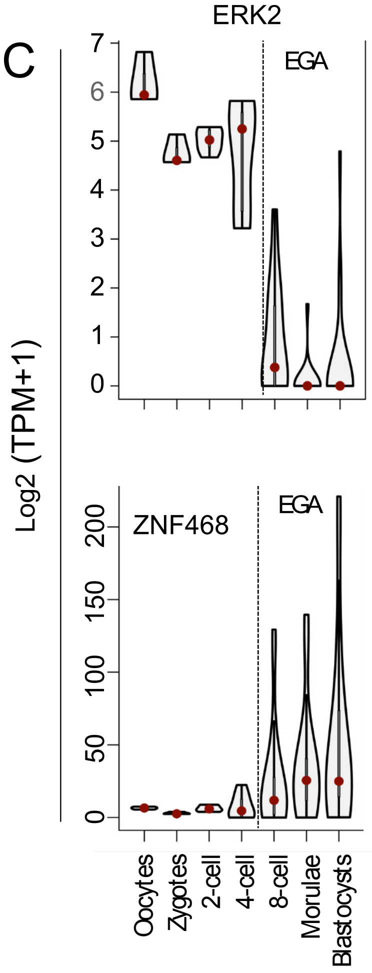


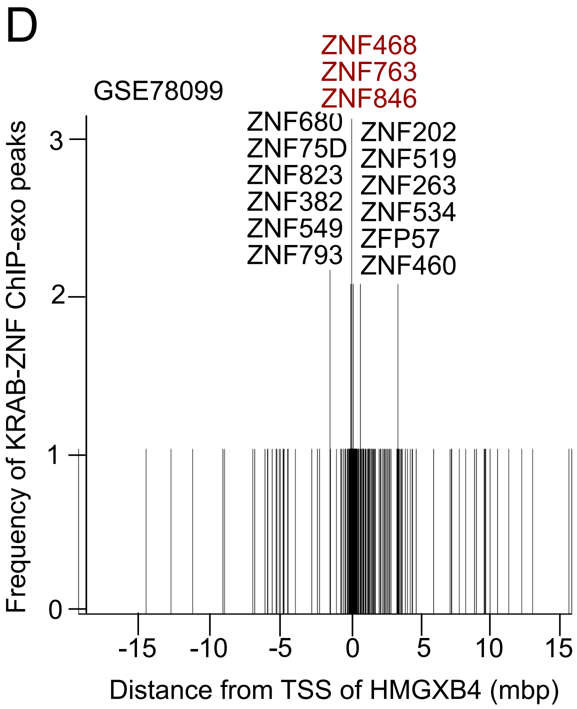


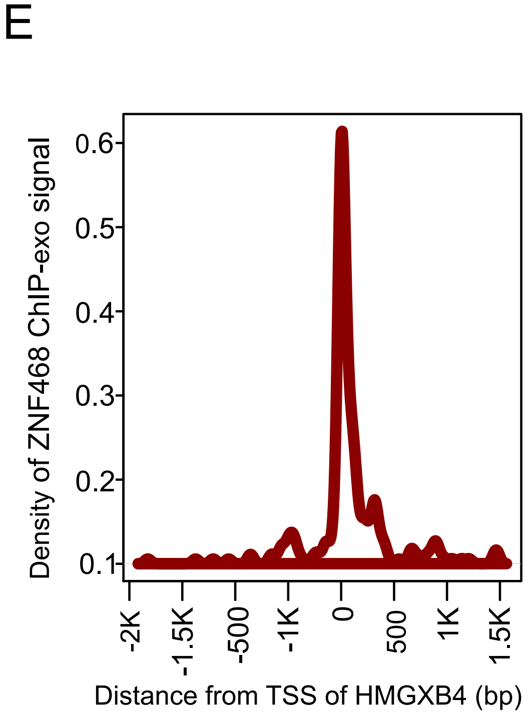


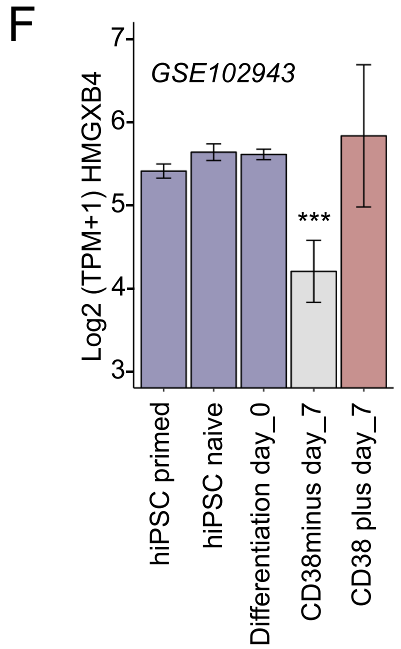


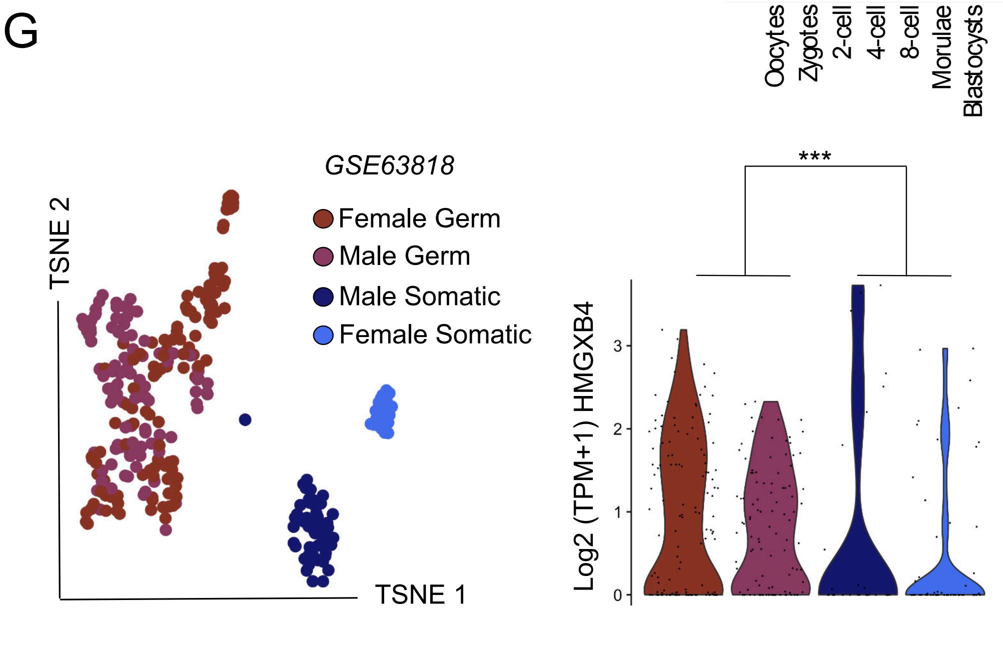
