## Supplementary figures and images for "HMGXB4 Targets *Sleeping Beauty* Transposition to Vertebrate Germinal Stem Cells"

### HMG Supplemental Figure 2

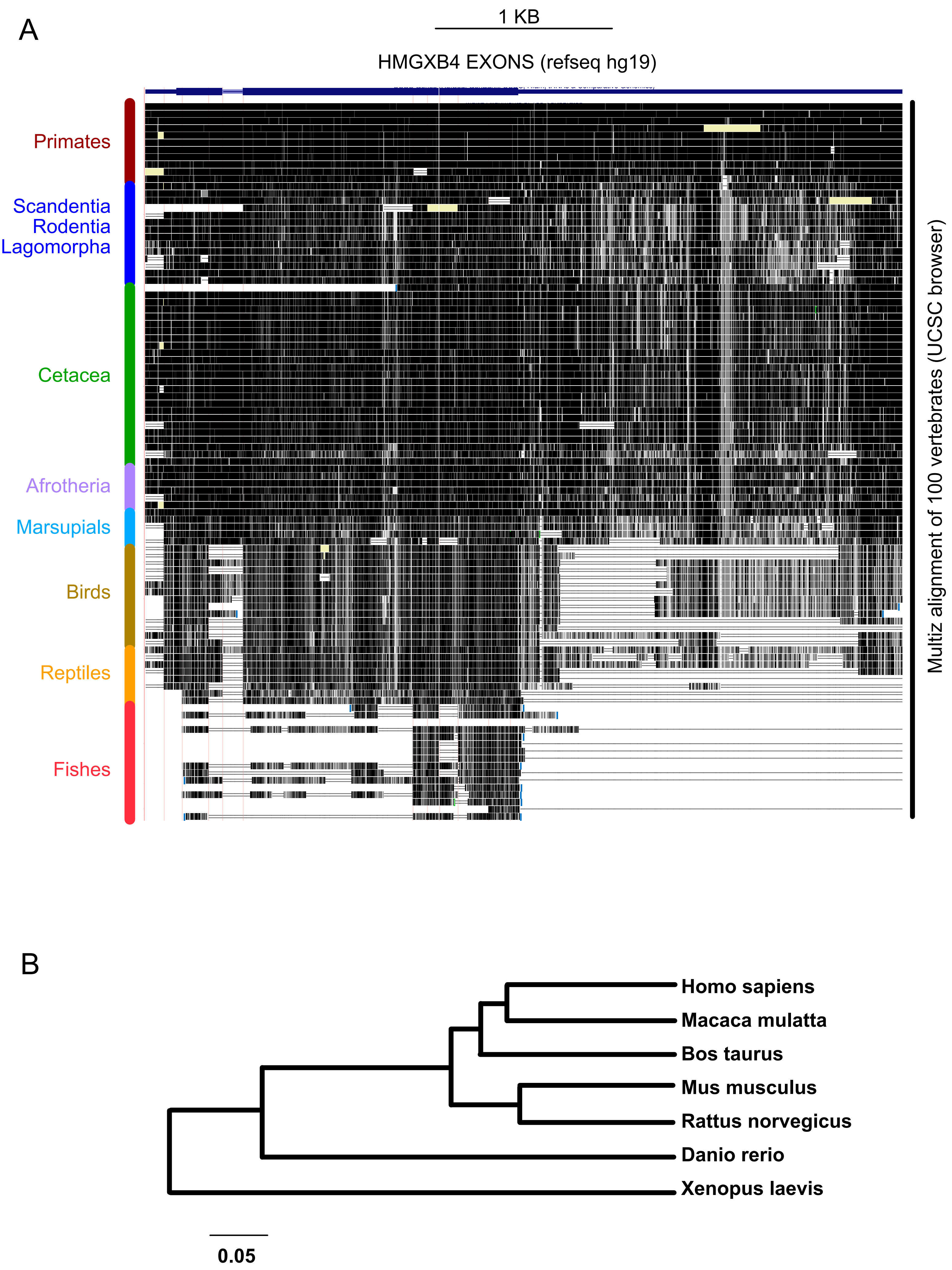


CLUSTAL-W alignment of HMGXB4 coding sequence

### HMG Supplemental Figure 3

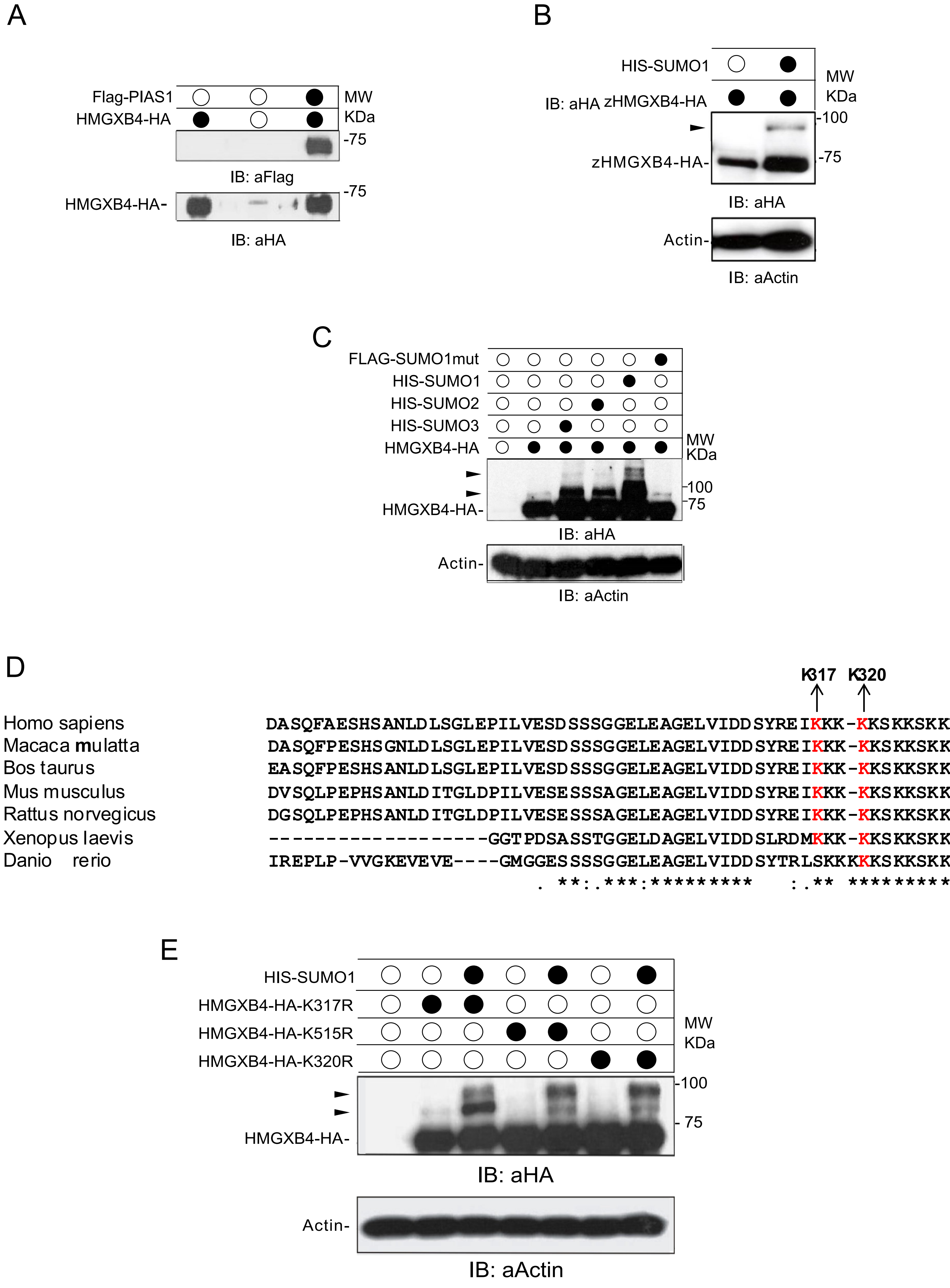

### HMG Supplemental Figure 5

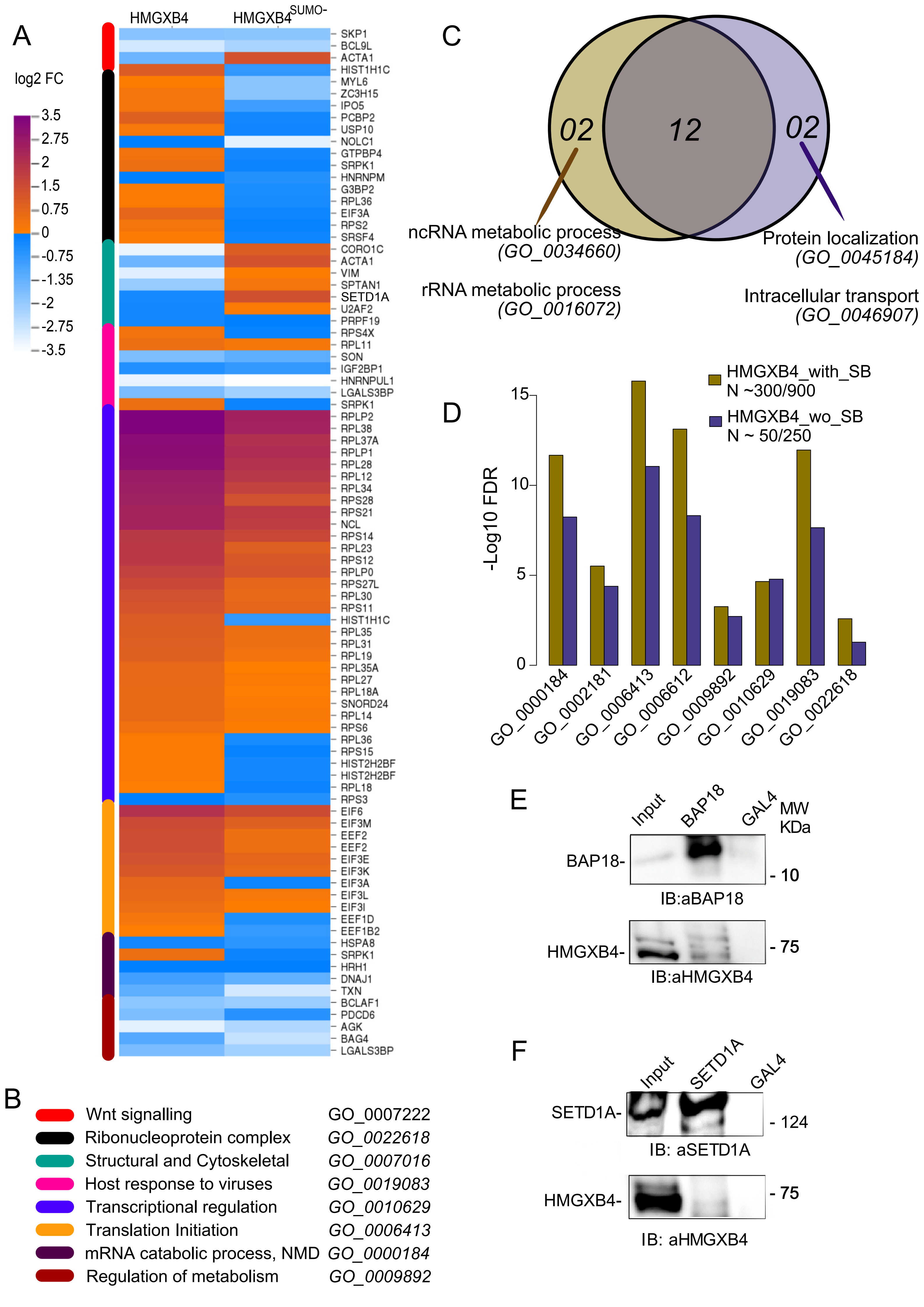

### HMG Supplemental Figure 6

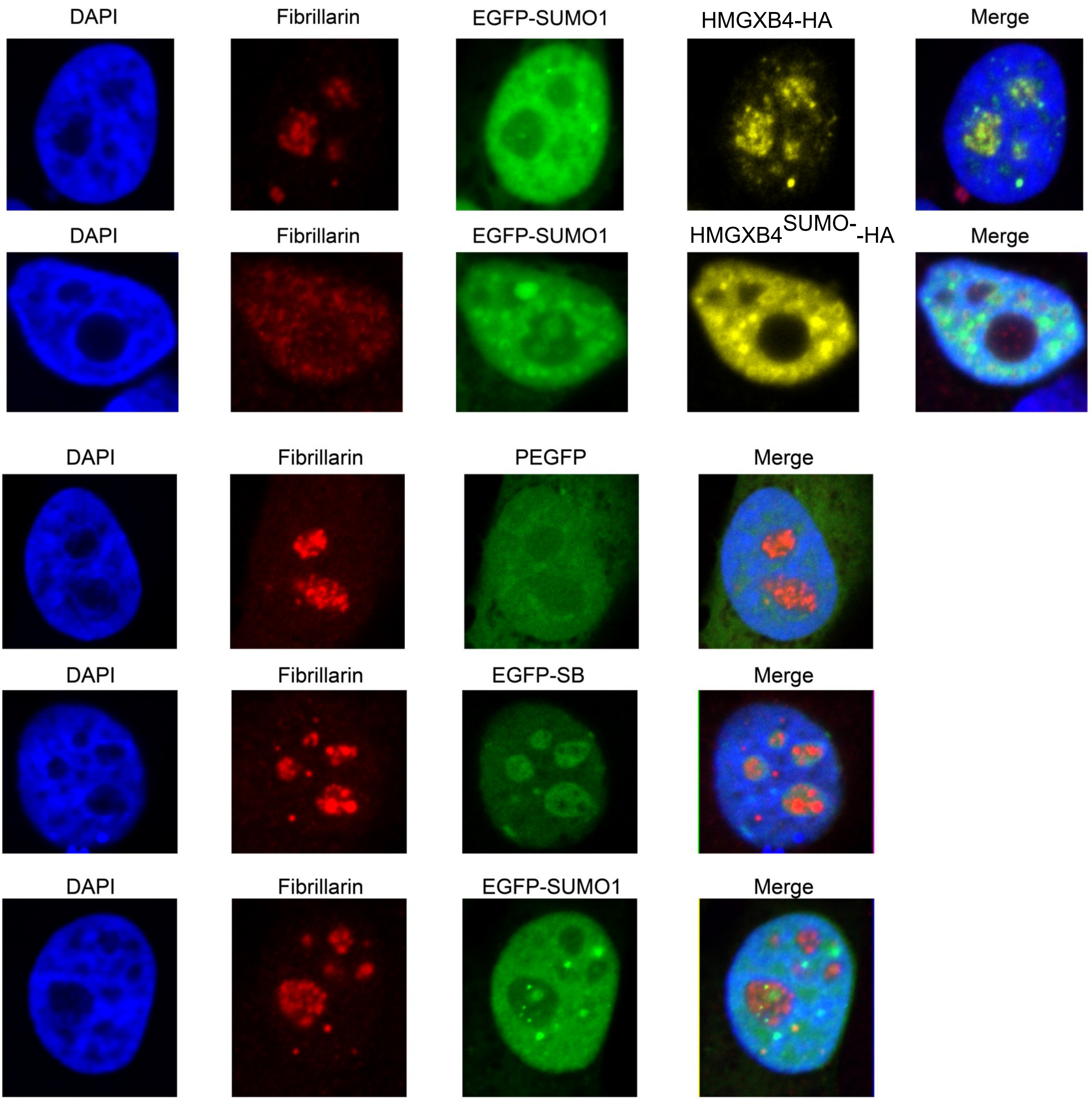
