## Supplementary material for "HMGXB4 Targets *Sleeping Beauty* Transposition to Vertebrate Germinal Stem Cells": HMG Supplemental Figure 4

A

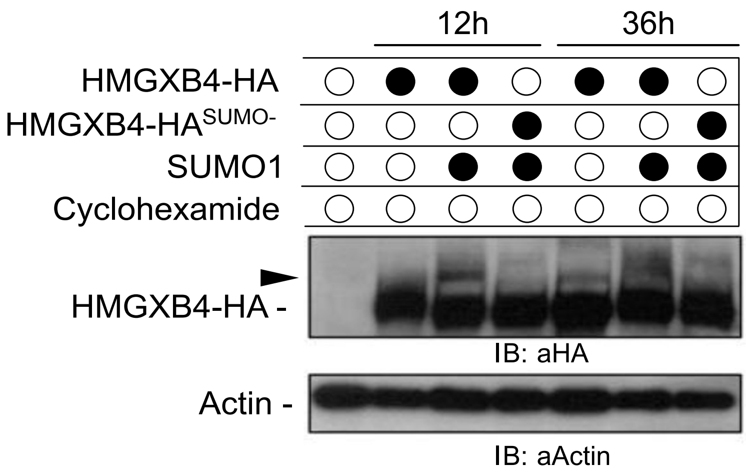


B

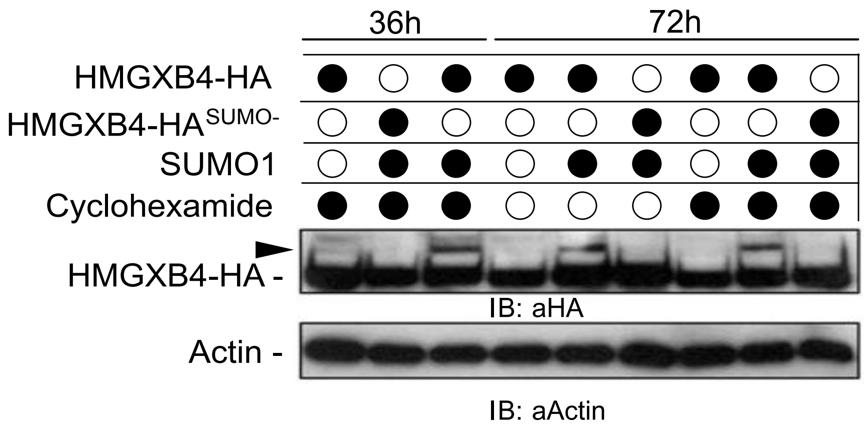


C

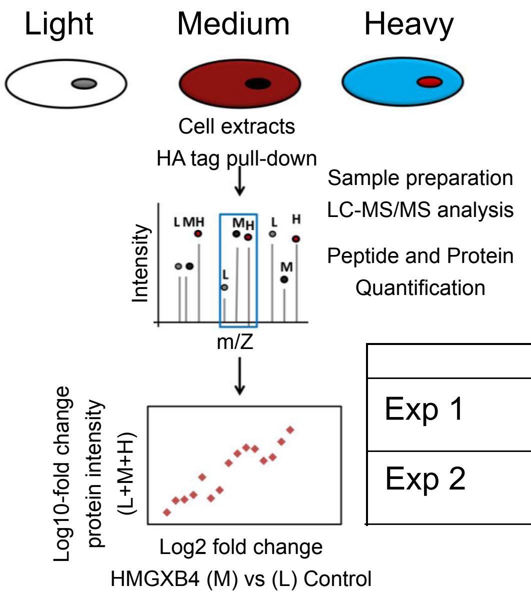


| | Light | Medium | Heavy |
|---|---|---|---|
| Exp 1 | HA only +SUMO1 | HMGXB4$^{SUMO-}$-HA +SUMO1 | HMGXB4 $^{WT}$-HA +SUMO1 |
| Exp 2 | HA only +SUMO1+SB | HMGXB4$^{SUMO-}$-HA +SUMO1+SB | HMGXB4 $^{WT}$-HA +SUMO1+SB |
